# Alizarin red perturbs skeletal patterning and biomineralization via Catalase inhibition

**DOI:** 10.1101/2025.07.03.663060

**Authors:** Abigail E. Descoteaux, Marko Radulovic, Mayank Ghogale, Santhan Chandragiri, Deema Abayawardena, Bikram D. Shrestha, Athula H. Wikramanayake, Vivek N. Prakash, Cynthia A. Bradham

## Abstract

Alizarin Red S (AZ) is an anthraquinone dye that is commonly used in histological studies and textiles. Exposure to AZ results in morphological perturbations in several species, including rats, frogs, and dogs; however, the mechanisms by which AZ has these effects is largely unexplored, and little is known about its effect on development. We have previously shown that AZ is teratogenic to sea urchin larvae, and that AZ was the only calcium-binding mineralization marker among five tested that perturbed skeletal patterning. Here, we further characterize these defects and demonstrate that embryos exposed to AZ have abnormal skeletal element rotation, branching, and bending. Immunostains and polychrome labeling reveal delayed migration of primary mesenchyme cells and initiation of biomineralization. Although gross ectodermal dorsal-ventral specification, ciliary band restriction, and neuronal specification occur normally in the majority of AZ-treated embryos, we find abnormal patterning and connectivity of the serotonergic neurons. Temporal transcriptomics comparisons confirm delayed development and implicate changes in neuron-related GO terms with AZ treatment. Particle imaging velocimetry experiments show that ciliary beating normally directs fluid flow into the larval mouth, while AZ treatment perturbs the normal pattern of vortices and redirects flow away from the mouth. These changes in fluid flow have functional consequences for larval feeding behaviors in AZ-treated embryos. Finally, we show that the effects of AZ on skeletal patterning are largely due to its inhibition of catalase and subsequent elevation of reactive oxygen species. Specifically, catalase knockdown and transient hydrogen peroxide treatment are each sufficient to phenocopy the hallmark AZ-mediated skeletal patterning defects. This study is the first to define the teratogenic consequences of AZ exposure and show how its effects on catalase activity impact development, skeletal patterning, and biomineralization.

**Summary Statement:** Alizarin Red exposure produces teratogenic effects on skeletal and nervous systems primarily via catalase inhibition.

## Introduction

Alizarin Red S (AZ) is a multifunctional anthraquinone dye commonly used in geological and histological studies to stain calcium-rich materials (Puchtler et al., 1969; Green, 2001; Arora et al., 2022). AZ is also a key constituent of madder, which has been used in textile dyes and paintings since ancient times (Fitzhugh, 1997). While AZ is widely useful, researchers have reported negative effects such as bone deformities and high mortality rates following AZ exposure during juvenile development in species such as rats, frogs, and dogs (Hoyte, 1960; Adkins, 1965; Rubin and Bisk, 1969; Lampertsdörfer et al., 1991). We recently demonstrated that AZ exposure perturbs skeletal pattern formation in sea urchin larvae as well (Descoteaux et al., 2023); however, the mechanisms by which AZ provokes these defects are largely unexplored, and little is known about its effect on development. Understanding how AZ mediates its effects is critical: AZ crosses the placenta (Shimidzu, 1922) and thereby poses a potential teratogenic threat to developing mammalian embryos including humans. Given these findings and because AZ’s industrial use and disposal are poorly regulated, AZ accumulation in the marine environment is an under-appreciated concern (Brown, 1987; Pereira and Alves, 2012; Routoula and Patwardhan, 2020; Moorthy et al., 2022). Understanding the mechanism by which AZ produces developmental defects may offer insight into strategies to prevent its teratogenicity.

To explore these questions, we use the larva of the sea urchin *Lytechinus variegatus*. The relatively simple calcium carbonate larval skeleton of *L. variegatus* is secreted by primary mesenchyme cells (PMCs), which ingress into the blastocoel during early gastrulation and migrate to a stereotypic pattern in response to cues from the overlying ectoderm (von Ubisch, 1937; Ettensohn and McClay, 1986; Ettensohn, 1990; Armstrong et al., 1993; Malinda and Ettensohn, 1994; Hardin and Armstrong, 1997; Hodor and Ettensohn, 1998; Tan et al., 1998; Wilt et al., 2008; Piacentino et al., 2015; Piacentino et al., 2016a; Piacentino et al., 2016b; Descoteaux et al., 2023; Hawkins et al., 2023; Thomas et al., 2023). Many of these ectodermally-expressed patterning cues, including BMP5-8, 5-LOX, and proteoglycans are highly conserved and are relevant to human skeletal development (Hästbacka et al., 1994; Traianedes et al., 1998; Kere, 2006; Cottrell et al., 2013; Wu et al., 2016; Xu et al., 2023). Thus, studies focused on how perturbations in gene expression and disruption of signaling pathways lead to defects in skeletal patterning and biomineralization in sea urchin larvae are likely to provide insight into similar mechanisms in humans and other vertebrates.

We show that AZ treatment prior to and during gastrulation in *L. variegatus* larvae produces abnormalities in gene expression and in PMC, skeletal, and neuronal patterns. We also find that fluid flow around the larvae is affected, indicative of abnormal ciliary movement and thus perturbed neural function, and that this results in decreased feeding. Finally, we demonstrate that AZ treatment globally increases reactive oxygen species (ROS) in sea urchin embryos and that AZ-mediated defects can be phenocopied by catalase knockdown or transient H_2_O_2_ exposure, suggesting that the effects of AZ are at least partially due to excessive reactive oxidation species. Together, these results show that catalase inhibition or ROS production are sufficient to perturb skeletal patterning, and thereby define a mechanism for the effects of AZ.

## Results

### AZ exposure during gastrulation produces dramatic skeletal patterning defects

Alizarin Red S (AZ) is a dye with a fluorescent ring structure and two calcium-binding moieties (Fig. 1A). When sea urchin larvae are exposed to AZ during development, AZ selectively labels the calcium carbonate skeleton (Fig. 1B); however, at sufficiently high concentrations, AZ-treated embryos exhibit severe skeletal patterning defects (Descoteaux et al., 2023). To characterize these defects, we first conducted dose-response experiments. We found that at low concentrations of AZ (≤ 10 μM), the majority of embryos were control-like or had mild rotational defects (Fig. 1C, Fig. S1A- B). At 15 μM AZ, most embryos had at least mild skeletal patterning defects such as rotational defects or stunted skeletal elements (Fig. 1C, Fig. S1C). Embryos treated with ≥ 30 μM AZ had severe patterning defects such as missing or spurious elements and dramatic rotational defects (Fig. 1C, Fig. S1D-F). As the concentration of AZ increased, the embryos produced less skeleton (Fig. S1G-H) and the angle of rotation between the two halves of the skeleton increased (Fig. S1A-F, I). At doses ≥ 75 μM, AZ precipitated out of the culture media and was consumed by the embryos (Fig. S1H, inset).

**Figure 1.**
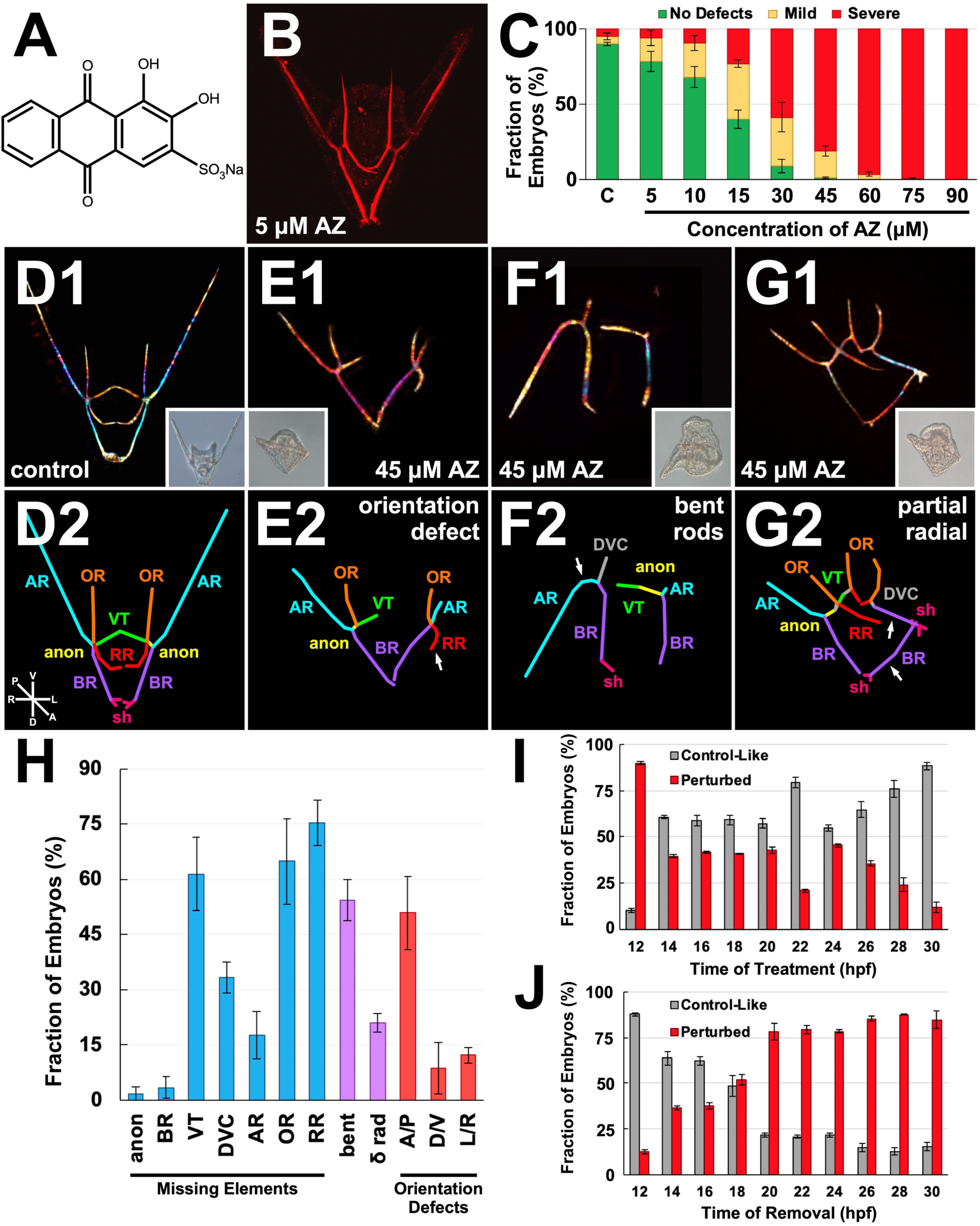
AZ treatment during gastrulation perturbs skeletal patterning in sea urchin embryos. **A.** The chemical structure of alizarin red S (AZ). **B.** Fluorescent signal of AZ in the larval skeleton. **C.** The fraction of embryos with normal (green), mildly perturbed (yellow), or severely perturbed (red) skeletal patterning after incubation in sea water (C) or the indicated concentrations of AZ (in μM) is displayed as the average percentage ± S.E.M. **D-G.** Skeletal patterns of control (D) and AZ-treated embryos (E- G) are shown at 48 hpf as skeletal birefringence (1) and schematically (2). Insets show corresponding morphologies (DIC). Skeletal elements in (2) are the anonymous rod (anon), ventral transverse rod (VT), dorsal-ventral connecting rod (DVC), body rod (BR), aboral rod (AR), oral rod (OR), recurrent rod (RR), and sheitel (sh). Embryonic orientation is indicated in D2. Arrows in E2-G2 indicate abnormally oriented, spurious, or atypically connected elements. **H.** The fractions of embryos with missing skeletal elements (blue), key morphologies (purple), and/or orientation defects (red) after treatment with 45 μM AZ are displayed as the average percentage ± S.E.M. The other scored defects are bent skeletal rods (bent); partially radialized (δ rad); and rotations about the anterior-posterior (A/P); dorsal-ventral (D/V); left-right (L/R). **I-J.** The fraction of embryos with normal (grey) or perturbed (red) skeletal patterning at 48 hpf after AZ is added (I) or removed (J) at the indicated time points during development is displayed as the average percentage ± S.E.M.

Because embryos treated with 45 μM AZ showed dramatic patterning defects without general inhibition of biomineralization or overt toxicity (such as gastrulation failures or excessive mesenchyme production), this dose was selected for further investigation. Compared to control larvae at similar time points (Fig. 1D), AZ-treated embryos showed dramatic rotational defects, especially about the anterior-posterior (A-P) axis, as well as missing skeletal elements (Fig. 1E-H). Aside from a few stereotypic locations, the skeletal elements of the pluteus-stage control larva are mostly straight, with branches forming at consistent angles and extending linearly along that plane (Fig. 1D) (Descoteaux et al., 2023); however, in AZ-treated embryos, many skeletal elements possess abnormal curvature, often bending at extreme angles relative to their initial direction of biomineralization after branching from the triradiates (Fig. 1E-F, arrows). A small but reproducible proportion of embryos (20%, n = 57) had a partially radialized phenotype, exhibited by a loss of the stereotypic bilateral symmetry and extra triradiates (Fig. 1G-H). Taken together, these data show that AZ treatment induces dramatic AP rotational defects, loss of skeletal elements, and bending of skeletal spicules, along with rarer instances of partial radialization.

To define *when* AZ exposure has these effects, we performed a series of timed treatment and removal experiments. For our timed treatment experiments, we exposed embryos to AZ every two hours from 12 to 30 hpf and then scored the resulting embryos at 48 hpf (Fig. 1I). We found that AZ treatment at or before 12 hpf is sufficient to perturb skeletal patterning with high penetrance in sea urchin embryos. Embryos remain partially sensitive throughout skeletal patterning until 24 hpf, after which their sensitivity declines (Fig. 1I). Next, to determine when AZ ceases to be effective, we performed timed removal experiments in which we added AZ at fertilization and then washed it out every two hours between 12 and 30 hpf. We found that a dramatic majority of embryos showed skeletal patterning defects when AZ was washed out as early as 20 hpf, while earlier removals resulted in mostly control-like embryos (Fig. 1J). Together, these experiments indicate that AZ is most effective at perturbing skeletal patterning between 12 and 20 hpf. This window correlates with gastrulation and the earliest biomineralization and patterning stages of the larval skeleton (Descoteaux et al., 2023).

### AZ treatment delays initiation of migration and perturbs triradiate orientation and branching

To better understand how skeletogenesis is affected by AZ treatment, we next performed polychrome labeling experiments (Descoteaux et al., 2023). Control and AZ-treated embryos were exposed to calcein green (CG) for only the first 24 hours of development, then imaged at 48 hpf. Since AZ-treated embryos also incorporate fluorescent label from AZ into the skeleton, we similarly exposed control embryos to xylenol orange (XO) for the duration of the experiment to rule out any differences due to the simultaneous presence of two fluorochromes during labeling. We found that, compared to control embryos (Fig. 2A), AZ-treated embryos incorporate less CG during the first 24 hours of development (Fig. 2B). This indicates that there is dramatically less biomineralization in AZ-treated embryos by 24 hpf, suggesting that initiation or elongation of biomineralization is disrupted in AZ-treated embryos.

**Figure 2.**
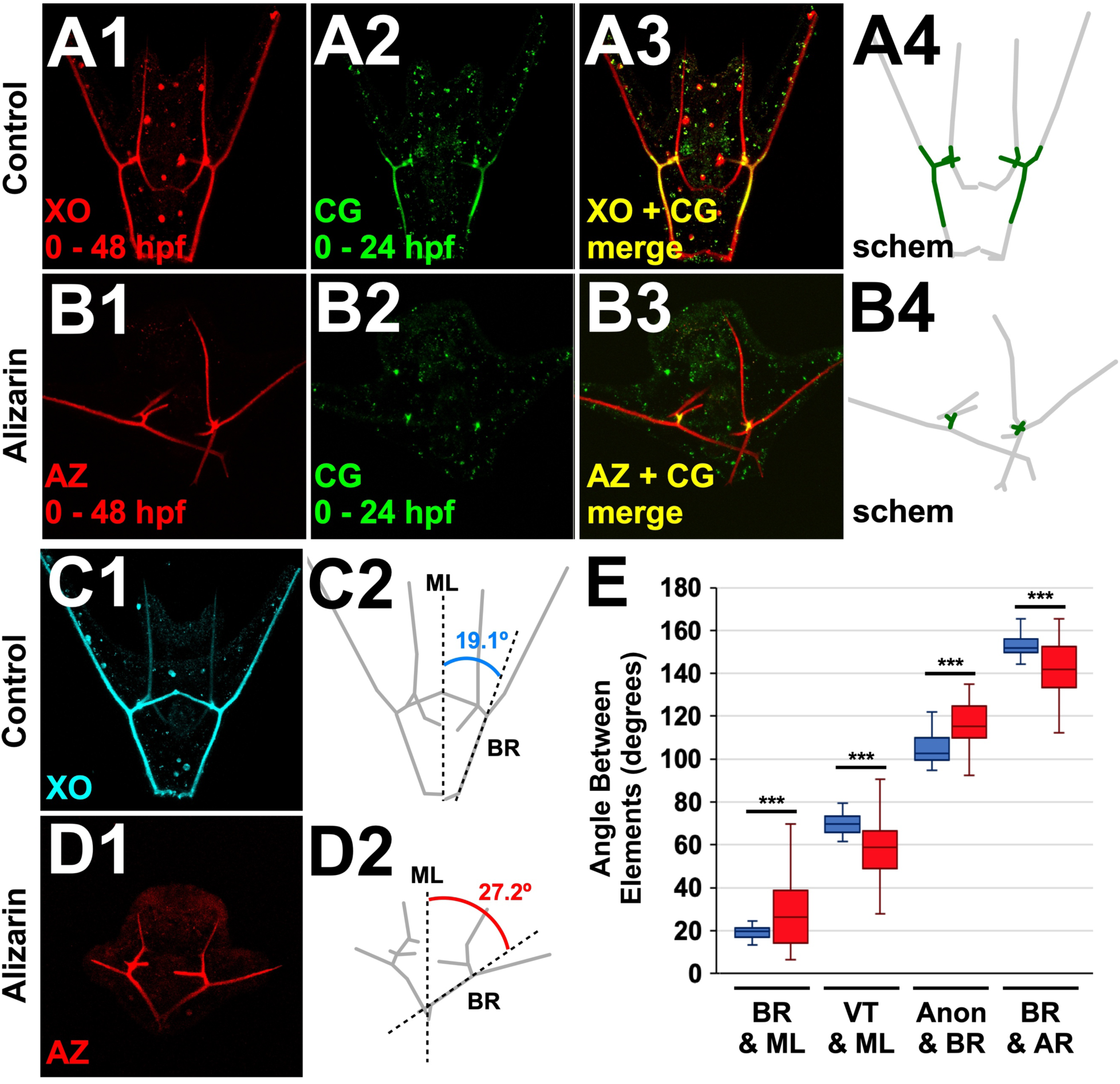
AZ treatment perturbs triradiate initiation, orientation, and branching. A-B. A control (A) and an alizarin-treated (B) embryo double-labeled with calcium-binding fluorochromes as indicated are shown as individual fluorochromes (1-2) and as a merge of calcein green (CG) with either xylenol orange (XO) (A3) or AZ (B3). The spatial extent of incorporation of the pulsed CG label (green) versus the entire skeleton (grey) is shown schematically (4). **C-D.** The average angle between the body rod (BR) and the midline (ML) was measured (2) in embryos whose full skeletons were labeled with XO (control, C) or AZ (D) at 48 hpf (1). **E.** The range of angle measurements between the indicated skeletal elements or to the ML are shown as box-and-whiskers plots, with the whiskers reflecting the 10^th^ and 90^th^ percentiles; n ≥ 17; *** p < 0.0005 (*t*-test). Skeletal elements are abbreviated as in Fig. 1.

AZ-treated embryos often exhibit dramatic A-P rotational defects, indicated by the abnormal positioning of the skeletal elements relative to the midline (Fig. 1E, H; Fig. 2B, D). To test if these rotational defects are due to overall rotation of the triradiates, abnormal branching of the skeletal elements, or a combination of these effects, we measured the angles in 3-D between several skeletal elements and the midline in control and AZ-treated embryos (Descoteaux et al., 2023). We found that all measured angles were significantly different in AZ-treated embryos (Fig. 2C-E, Fig. S2), indicating that the observed A-P rotational defects result from both abnormal rotation of the triradiates and structural changes in the branching angles, specifically the anonymous rod into the body rod and aboral rod. Taken together, these data suggest that AZ treatment perturbs not only the patterning of, but also the biomineralization of the skeletal elements.

### PMC patterning and identity are disrupted in AZ-treated embryos

In sea urchin larvae, the skeletal biomineral is secreted by the PMCs into the lumen of their shared syncytial cable; thus, the spatial arrangement of the PMCs dictates the pattern of the ensuing skeleton. Since we observed dramatic defects in the skeletal pattern of AZ-treated embryos, we next investigated how the migration and spatial positioning of these skeletogenic cells is affected by AZ treatment via immunolabeling PMCs at 24 and 48 hpf (Fig. 3A-B). We found that at 24 hpf, the migration of the PMCs in AZ-treated embryos was dramatically delayed (Fig. 3A1, B1), with PMCs still in the ring-and-cords pattern characteristic of 18 hpf control embryos (Adomako-Ankomah and Ettensohn, 2013; Hawkins et al., 2023; Lyons et al., 2012; Piacentino et al., 2016a; Rodríguez-Sastre et al., 2023; Thomas et al., 2023; Zuch and Bradham, 2019). This delay is consistent with our polychrome labeling findings (Fig. 2A- B). At 48 hpf, a significant fraction of AZ-treated embryos had defects in PMC positioning, such as ectopically located PMCs and clusters (Fig. 3B2 and Fig. S3C, arrows), abnormal webbing of the PMC filopodia (Fig. S3B, arrows), and stalled plugs of PMCs (Fig. 3C). These data show that AZ treatment both delays and disrupts PMC migration and patterning.

**Figure 3.**
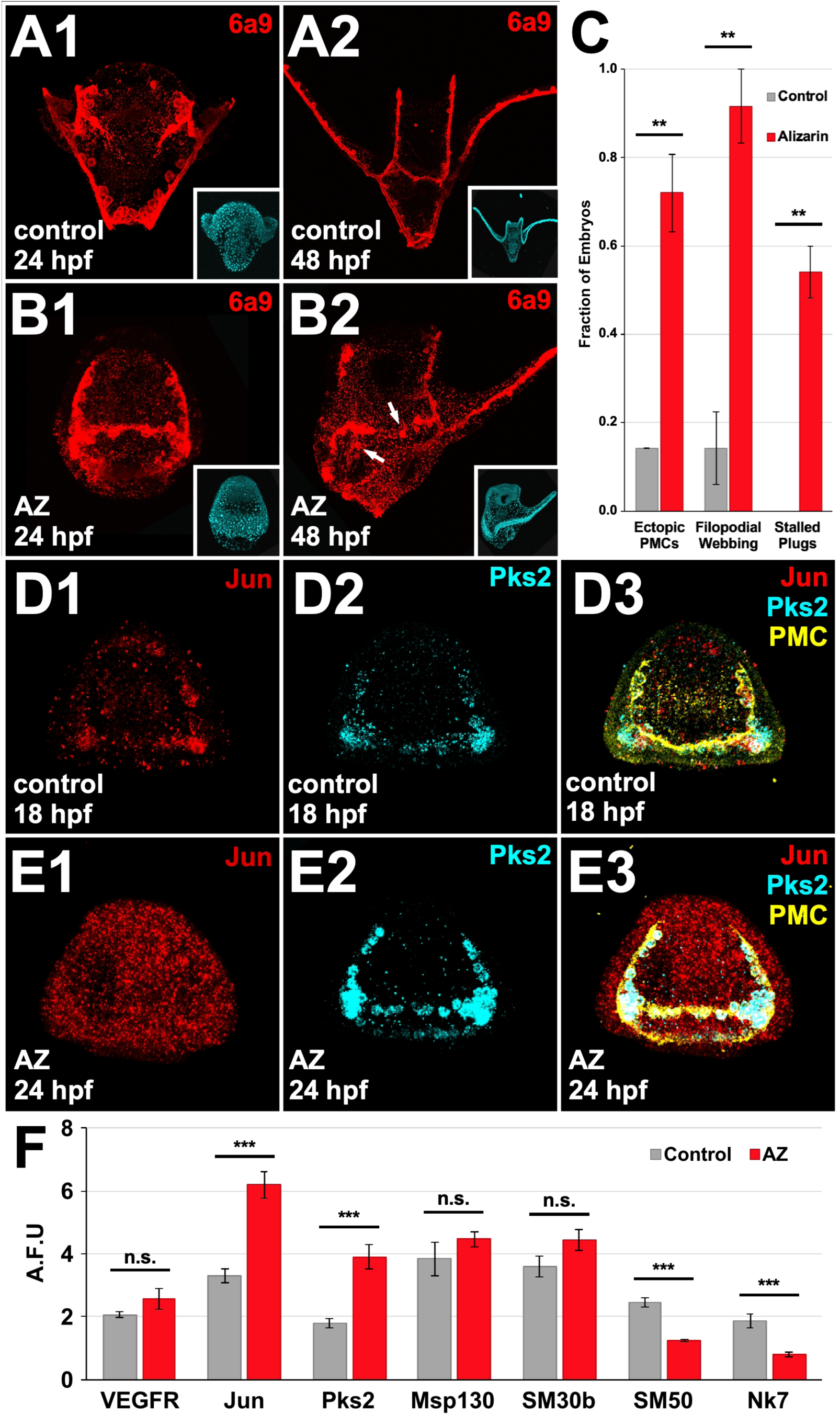
AZ treatment perturbs PMC migration, patterning, and identity. A-B. Immunolabeled PMCs in control (A) and AZ-treated embryos (B) are shown at 24 (1) and 48 hpf (2). Insets show nuclei labeled with Hoechst in the corresponding embryo. Arrows in B2 indicate ectopic PMCs. **C.** The fraction of embryos at 48 hpf showing the indicated defect in control (grey) and AZ-treated embryos (red) is shown as average percentage ± S.E.M; n ≥ 21; ** p < 0.005 (*t*-test). **D-E.** Stage-matched control (D) and AZ-treated (E) embryos were subjected to HCR FISH for jun (red) and pks2 (cyan) followed by PMC immunolabeling (yellow). FISH signals are shown individually (1-2) and merged with PMC stain (3). **F.** The normalized average expression per unit area of the indicated PMC subset genes is shown as the average artificial fluorescent unit (A.F.U.) ± S.E.M in control (grey) and AZ-treated embryos (red); n ≥ 15; *** p < 0.0005; n.s. not significant (*t*-test).

As the PMCs migrate during skeletal patterning, they also diversify into spatially-defined subsets with unique gene expression profiles (Armstrong et al., 1993; Duloquin et al., 2007; Adomako-Ankomah and Ettensohn, 2013; Sun and Ettensohn, 2014; Piacentino et al., 2015; Piacentino et al., 2016a; Zuch and Bradham, 2019; Hawkins et al., 2023) using stage-matched comparisons of controls at 18 hpf and AZ-exposed embryos at 24 hpf. We tested whether PMC identity was affected by AZ treatment using single molecule fluorescent in situ hybridization (smFISH) (Choi et al., 2018; Choi et al., 2020) to look for changes in the expression levels or spatial domains of several PMC subset genes. A well-studied PMC subset gene is VEGFR, the receptor for the ectodermal cue vascular endothelial growth factor (VEGF) (Duloquin et al., 2007; Röttinger et al., 2008; Adomako-Ankomah and Ettensohn, 2013; Piacentino et al., 2016a). However, we found no significant difference in *vegfr* expression in embryos treated with AZ (Fig. 3F, Fig. S4A); consistent with this outcome, the AZ phenotypes are dissimilar to those of late VEGFR inhibition (Adomako-Ankomah and Ettensohn, 2013; Piacentino et al., 2016a) suggesting that this signaling pathway is not involved in mediating the effects of AZ on skeletal patterning. Another cluster-specific PMC subset gene is the transcription factor Jun, which is normally initially globally expressed before being restricted to the PMC clusters (Rodríguez-Sastre et al., 2023; Sun and Ettensohn, 2014); however, in AZ-treated embryos, *jun* loses this spatial restriction and remains globally elevated (Fig. 3D1, E1, F). Polyketide synthase 2 (*pks2*), which is normally highly expressed in the PMCs clusters and only weakly expressed by other PMCs, (Castoe et al., 2007; Hawkins et al., 2023; Sun and Ettensohn, 2014; Zuch and Bradham, 2019), is significantly elevated in the PMCs of AZ-treated embryos (Fig. 3D2, E2, F). Other PMC subset genes include those encoding Msp130, SM30b, and SM50, which are biomineralization and spicule matrix proteins, and Nk7, a homeobox transcription factor (Wilt et al., 2008; Killian et al., 2010; Mann et al., 2010; Rao et al., 2013). mRNAs encoding Msp130 and SM30b are not significantly differently expressed in AZ-treated embryos; however, SM50 and Nk7 are both significantly decreased (Fig. 3F, Fig. S4B-E). Thus, we find that AZ treatment alters the expression of some PMC subset genes, suggesting some loss of PMC identities in AZ-treated embryos.

### AZ treatment does not affect the expression of Univin, Wnt5, or VEGF at late gastrula stage

The migration and patterning of the PMCs is instructed by cues from the overlying ectoderm (von Ubisch, 1937; Ettensohn and McClay, 1986; Ettensohn, 1990; Armstrong et al., 1993; Malinda and Ettensohn, 1994; Hardin and Armstrong, 1997; Piacentino et al., 2015; Piacentino et al., 2016a; Piacentino et al., 2016b; Descoteaux et al., 2023; Hawkins et al., 2023; Thomas et al., 2023). The PMC abnormalities that we observed in AZ-treated embryos could reflect perturbed ectodermal expression of patterning cues. VEGF, Univin, and Wnt5 are critical for normal PMC migration, while VEGF and Wnt5 are also required for biomineralization (Duloquin et al., 2007; Adomako-Ankomah and Ettensohn, 2013; McIntyre et al., 2013; Piacentino et al., 2015; Piacentino et al., 2016a; Thomas et al., 2023). To test if the expression of these patterning cues is affected by AZ treatment, we performed FISH in late gastrula-stage control and AZ-treated embryos. We found no significant difference in the expression domain or the relative expression level of these ectodermal genes in AZ-treated embryos (Fig. S5), suggesting that the abnormalities in PMC migration and identity that we observed in AZ-treated embryos are not due to disruption of these cues.

### AZ treatment delays and disrupts gene expression

To further understand how AZ treatment affects gene expression, we examined the transcriptomes of control and AZ-treated embryos at several time points during development using Illumina sequencing. We assessed the overall variation in the data using principle component analysis (PCA) (Fig. 4A) (Blighe et al., 2024). The transcriptome data was sorted by time along PC1, which is typical for PCA of sea urchin transcriptome data (Hogan et al., 2020; Wygoda et al., 2014). Interestingly, we see that control and AZ samples are similarly positioned along PC1 at 12 and 18 hpf but that AZ samples then exhibit a temporal delay such that 24 hpf AZ samples cluster more closely to 18 hpf control samples and 30 hpf AZ samples cluster more closely to 24 hpf control samples than to their respective time-matched samples. This agrees with our earlier observations of delayed development at 24 hpf in AZ-treated embryos (Fig. 2; Fig. 3).

**Figure 4.**
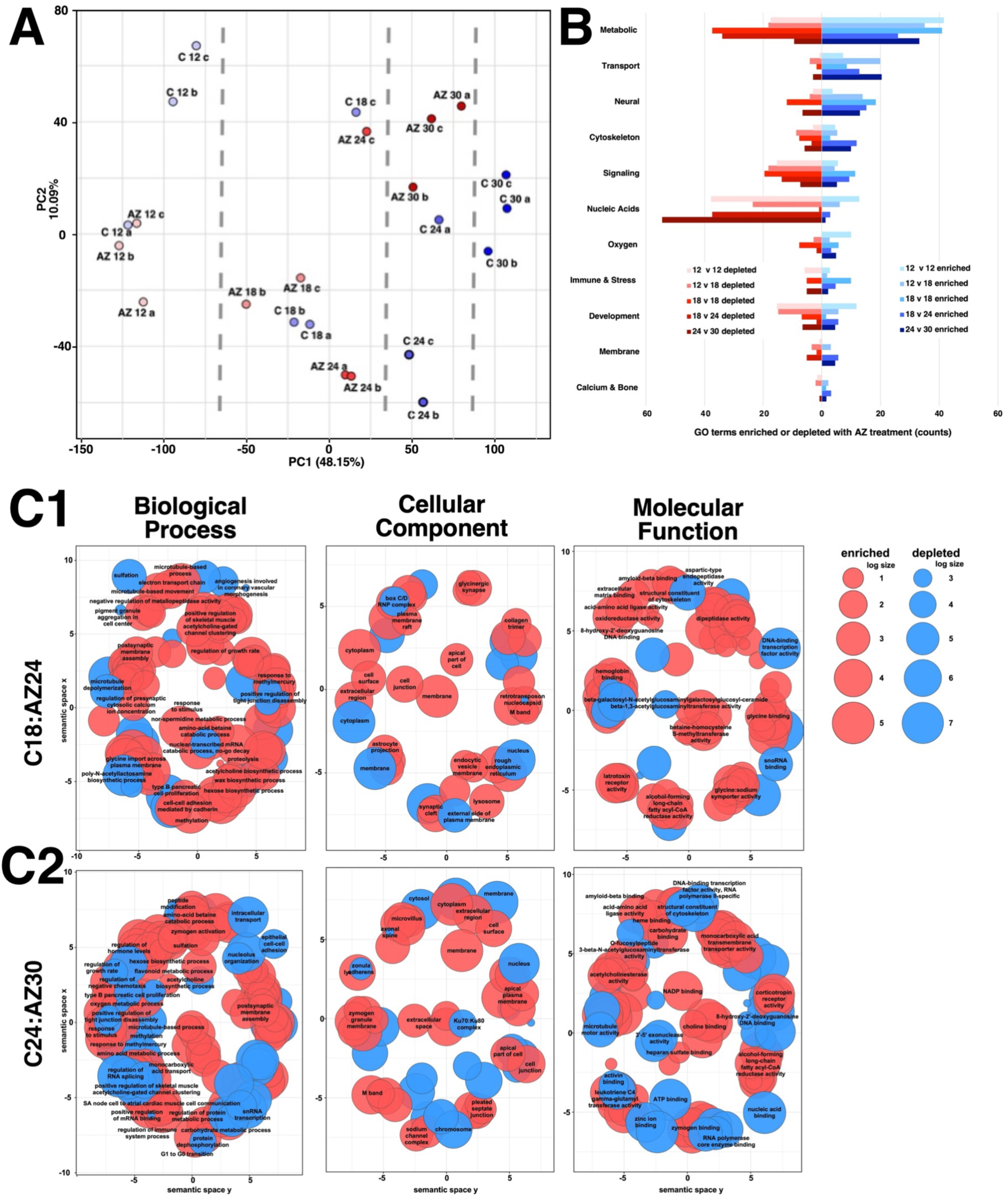
AZ treatment leads to delayed development and broad changes in gene expression. **A.** PCA is shown for the first two components for control (C) and AZ- treated (AZ) samples. Vertical dotted lines subdivide the plot by control time points along PC1. **B.** GO term enrichment for control vs AZ in the indicated homo- and heterochronic pairs is shown after binning into the indicated categories. See also Table S1. **C.** Semantic clustering of GO terms enriched and depleted by AZ is shown for two heterochronic comparisons; see also Fig. S6.

To examine how gene expression is broadly affected by AZ treatment, we analyzed the relative enrichment of gene ontology (GO) terms within several categories (Fig. 4B, Table S1). Because we observed temporal delay in gene expression in AZ samples, we performed this analysis for both time-matched and heterochronic comparisons. We find that the largest category affected by AZ treatment is metabolism, with slightly more GO term enrichment than depletion. Nucleic acids, which includes transcription, translation, splicing, recombination, and DNA repair, was the most highly depleted category in AZ samples (Table S1). Signaling and development categories were also strongly depleted in AZ, while transport and neural categories were each enriched (Fig. 4B). Oxygen-related GO terms, such as oxidoreductase activity and response to oxidative stress, were also slightly more enriched in AZ at most timepoints with the exception of 18 hpf, when there was a relative depletion of these terms.

For more granular insights into these effects, we analyzed the enriched and depleted GO terms using semantic clustering (Supek et al., 2011). These results reveal widespread effects throughout the cell, with broad metabolic and neural development impacts, along with smaller effects on development and morphogenesis, transport, cytoskeletal regulation, and adhesion (Fig. 4C, Fig. S6). Biological processes are most strongly impacted, while cellular components are least affected. However, specific pathways or other overarching connections between the affected GO terms did not emerge from these analyses. Together, these results thus suggest that AZ treatment leads to a wide range of gene expression changes across numerous gene categories and cellular compartments, with the largest impacts on genes associated with metabolism, nucleic acid regulation, transport, signaling, and neural development.

### AZ treatment does not disrupt ectodermal dorsal-ventral specification

A conspicuous ectodermal feature in sea urchin larva is the ciliary band: a narrow strip of ciliated cells whose beating powers larval swimming and feeding. The ciliary band is specified as the default non-neural ectodermal state, then is spatially restricted to the dorsal-ventral (DV) boundary via TGFβ-mediated specification of the DV regions (Bradham et al., 2009; Yaguchi et al., 2010). The spatial restriction of the ciliary band can therefore be used as a readout for whether ectodermal dorsal and ventral specification occurred normally (Piacentino et al., 2016a). We examined ciliary band specification in AZ-treated embryos via immunolabeling and found that in most AZ-treated embryos, the ciliated structures were normally restricted (80%, n = 30) (Fig. 5A- B); however, a small fraction of AZ-treated embryos displayed localized abnormalities in which ciliated band restriction was compromised (20%, n = 30) (Fig. 5C), implicating abnormal specification of the dorsal and/or ventral ectoderm.

**Figure 5.**
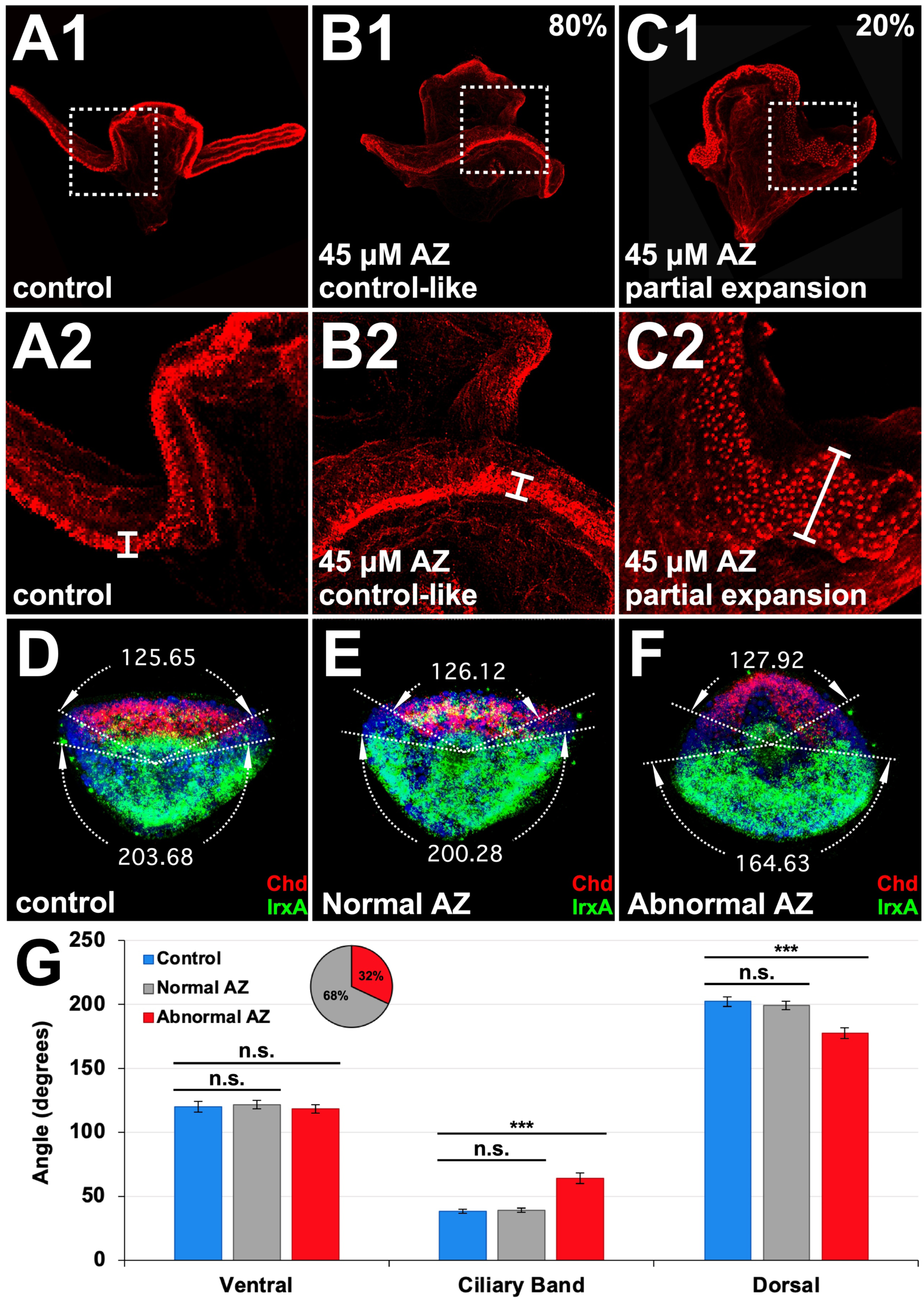
AZ treatment affects the ciliary band and ectodermal dorsal-ventral specification in a minority of embryos. A-C. Control (A) and AZ-treated embryos (B- C) were immunolabeled at 48 hpf to visualize the ciliary band. AZ-treated embryos show both normal restriction (B) and partial expansion (C) of the ciliary band, at the indicated frequencies. Whole embryos (1) and magnified views (2) of the indicated regions are shown. White bars in (2) highlight the width of the ciliary band. **D-F.** Control (D) and AZ- treated embryos (E-F) were subjected to HCR FISH for chordin (Chd, red) and IrxA (green) at late gastrula stage to measure the angular extent of the ventral and dorsal ectodermal regions. **G.** The measured angles of the ventral and dorsal ectodermal regions are shown as average angle ± S.E.M. The area between the Chd and IrxA expression domains was inferred as the presumptive ciliary band; *** p < 0.0005; n.s. not significant (*t*-test). Inset pie chart shows the fraction of AZ-treated embryos with normal (grey) or abnormal (red) inferred ciliary band area; n = 25.

To assess this further, we used FISH for dorsal and ventral marker genes *irxA* and chordin, respectively, to quantify the radial extent of the dorsal and ventral regions at late gastrula stage (Thomas et al., 2023). Since embryos at this stage are approximately circular when viewed from a vegetal perspective, the relative angular size of the presumptive ciliary band can also be inferred from these measurements. In control embryos, ventral *chordin* occupies approximately 85° of the embryo and dorsal *irxA* occupies approximately 220°, meaning that the two presumptive ciliary band regions combined occupy the remaining 55°. The majority of AZ-treated embryos had normal restriction of *chordin*, *irxA*, and the ciliary band (Fig. 5D-E, G). In the remaining AZ-treated embryos, although the spatial extent of the ventral *chd* domain was not significantly different than controls, the extent of the dorsal *irxA* expression domain was significantly smaller, resulting in a larger calculated angle of the combined ciliary band regions (Fig. 5F-G). These data suggest that in this fraction of AZ-treated embryos, the ciliary band is expanded due to a reduction of the dorsal ectoderm territory. These findings match the immunolabeling results above, along with the previously observed partial skeletal radialization in some AZ-exposed larvae.

### AZ treatment perturbs serotonergic neuron patterning and connectivity but not gross neuronal specification

The neurons of the sea urchin larvae are derived from the ectoderm; thus, their development and location are also indicators of DV specification. To learn whether neuronal specification was affected by AZ treatment, we first assessed global neuronal specification via FISH for the pan-neural gene synaptotagmin B (synB). We found that neurons were specified in similar locations and that the total number of neurons was comparable in AZ-treated embryos compared to controls (Fig. 6A-C). We also examined serotonergic (ser) neurons, which are present in the oral hood and are thought to be the central nervous system of the larva (Bisgrove and Burke, 1986; Yaguchi et al., 2006; Angerer et al., 2011), at 48 hpf via immunolabeling. We found no significant difference in the location or number of ser neurons in AZ-treated embryos (Fig. 6D-F); however, the ser neurons were often further apart and were lacking projections in AZ-treated embryos (Fig. 6E2, arrows). Taken together, these findings show that although neuronal specification occurs normally in AZ-treated embryos, the connectivity of the ser neurons is abnormal.

**Figure 6.**
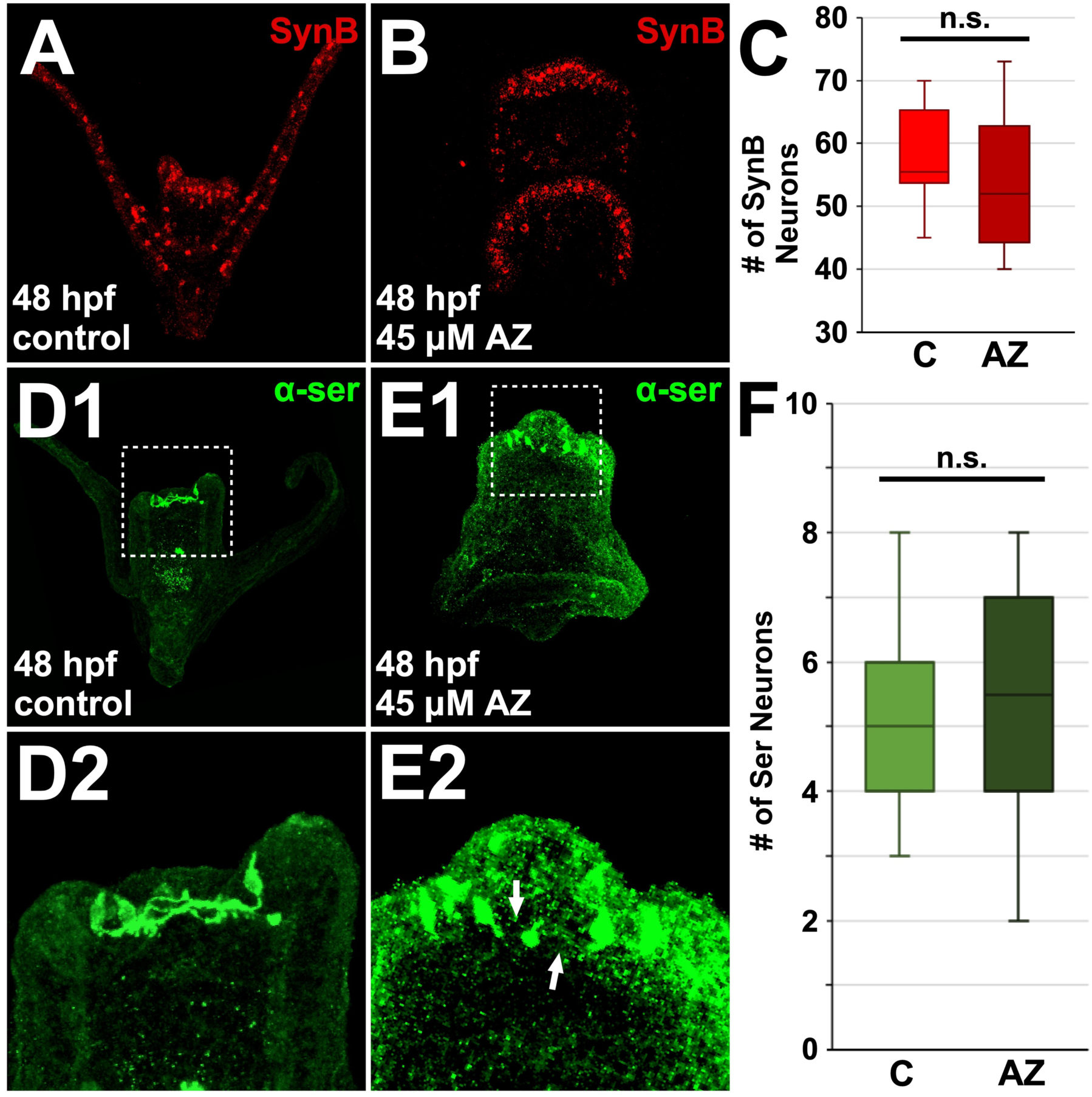
AZ treatment mildly perturbs serotonergic neuronal patterning but not overall neuronal development. A-B. Control (A) and AZ-treated embryos (B) at 48 hpf were subjected to HCR FISH for synaptotagmin B (SynB, red) to label all neurons. **C.** Average numbers of neurons in control (C) and AZ-treated (AZ) embryos at 48 hpf are shown as box-and-whiskers plots, with the whiskers reflecting the 10^th^ and 90^th^ percentiles; n ≥ 14; n.s. not significant (*t*-test). **D-E.** Control (D) and AZ-treated embryos (E) at 48 hpf were immunolabeled within anti-serotonin antibody (α-ser, green) to show distribution of serotonergic neurons. Whole embryo views (1) and magnified views of the oral hood (2) are shown. Arrows in E2 indicate missing neuronal projections. **F.** Average numbers of serotonergic (ser) neurons in control and AZ-treated embryos at 48 hpf are shown as box-and-whiskers plots, as in C.

### AZ-treated embryos produce abnormal fluid flow patterns

The beating of the cilia within the ciliary band is regulated by neurons (Mackie et al., 1969; Satterlie and Cameron, 1985). Although ciliary band restriction is normal in most AZ-treated embryos, the overall shape of the ciliary band is abnormal due to the abnormal morphologies of the AZ-treated embryos. Since the beating of the cilia within the ciliary band directs fluid flow around the larvae to facilitate swimming and feeding, we were curious if the abnormal morphology and abnormal connectivity of ser neurons in AZ-treated embryos would have consequences for fluid movements around the embryos. To examine this, we visualized the fluid flow patterns generated by ciliary beating by visualizing the movement of beads around live larvae under confinement (Shrestha et al., 2025) utilizing Flowtrace (Gilpin et al., 2017) and the Particle Image Velocimetry (PIV) technique (Thielicke and Stamhuis, 2014) for data post-processing and flow quantification (Fig. 7). We found that control and AZ-treated embryos each produced regions with rotational flows, or vortices, but that the number, location, and strength of these vortices varied with treatment. In confined control embryos, ciliary movements produced two to three pairs of vortices, with one pair near the dorsal end of the embryo and additional pairs on the medial and lateral side of the distal aboral arms (Fig. 7A1, D). Overall flow velocity is higher ventrally, near the mouth, than dorsally (Fig. 7A2). The ventrally increased velocity and vorticity of flow likely facilitates feeding by directing food particles in the water into the mouth. The intensity and directionality of fluid rotation, or vorticity (Fig. 7A3-4) alternates between negative vorticity/clockwise flow (blue) and positive vorticity/counterclockwise flow (red) in a manner that directs the overall fluid flow along the body from ventral to dorsal; interestingly, we find that the location and directionality of the vortices around the ventral region of the embryo result in flow of water towards the mouth of the embryo (Fig. 7A4).

**Figure 7.**
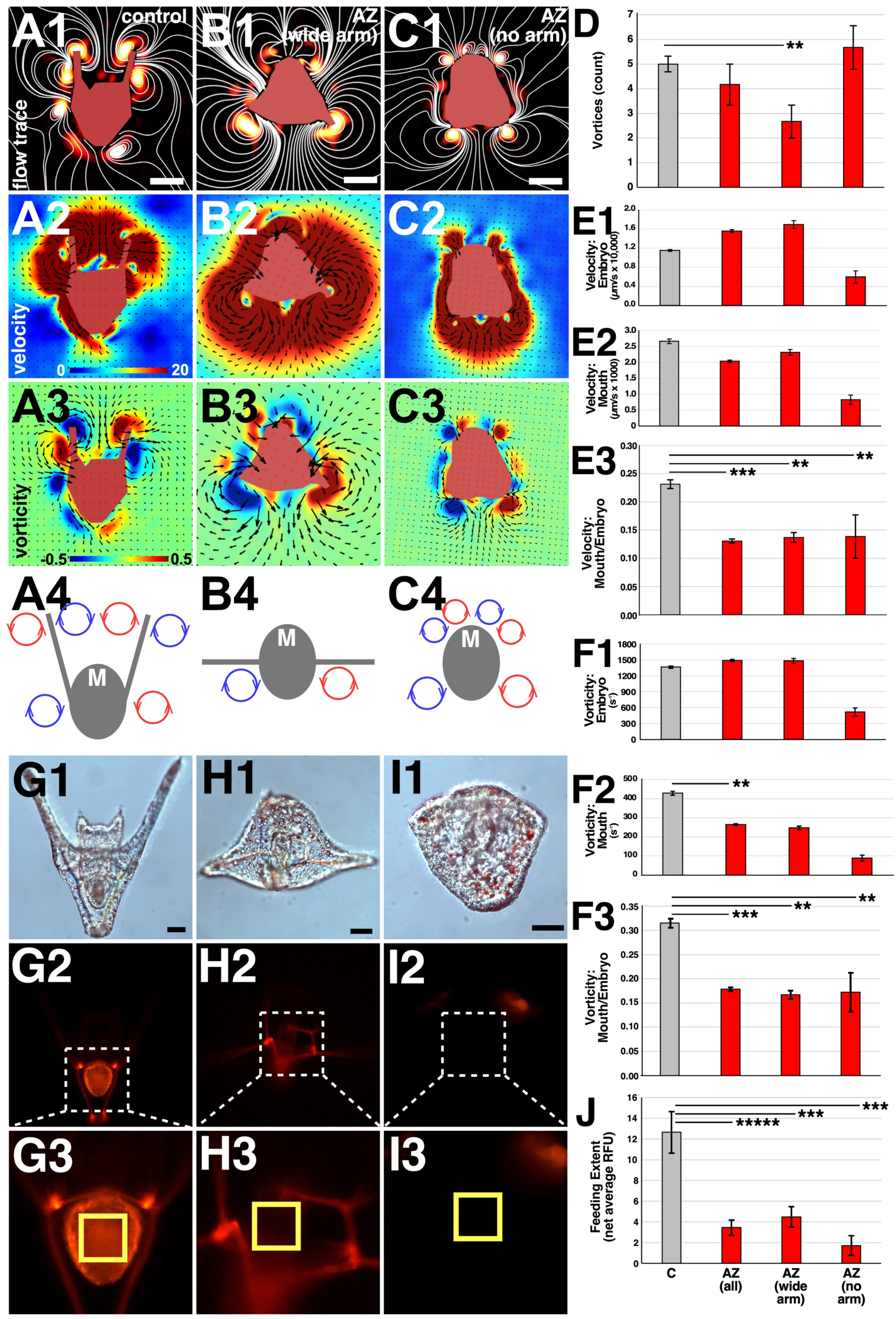
AZ treatment affects fluid flow around sea urchin larvae. A-C. Exemplar flow fields averaged across three seconds for control (A) and AZ-treated embryos (B-C) are shown as streamlines (1, white lines), magnitude of velocity (2), vorticity (3), and schematically (4). Embryos in 1-3 are masked in red and flow-fields are colorized with the indicated inset LUTs (A2, A3). Black lines in 2 and 3 indicate the vector fields. The mouth (M) of each embryo is also indicated in 4. **D**. The number of vortices is shown as the average ± S.E.M. **E-F**. The velocity (E) or vorticity (F) magnitude around the whole embryo (1), near the mouth (2), and the ratio of the magnitude near the mouth to the whole embryo (3) are shown as the average ± S.E.M. N = 5 for control embryos; n = 3 for AZ wide arms phenotypes; and n = 3 for AZ no arms phenotypes. **G-I**. Feeding experimental results with controls (G), wide arm AZ embryos (H) and no arm AZ embryos (I) are shown as morphology (DIC, 1), and as fluorescence of the whole embryo (2) and zoomed on the gut as indicated (3); the region for quantification is boxed in yellow. Scale bar corresponds to 50 µm. Linear and large spots of fluorescent signals outside the gut correspond to skeletal labeling by AZ (or XO in controls, see methods) rather than fluorescence from ingested algae and are not relevant to this experiment. **J.** Gut-specific fluorescence is shown as the average net relative fluorescence units (RFUs) + S.E.M. N = 35 for controls, 25 for AZ wide arms, and 15 for AZ no arms. ** 0.05 < p < 0.01; *** 0.01 > p > 0.001; ***** 10^-5^ < p < 10^-4^; otherwise not significant (*t*-test).

To quantify these effects, we focused on the velocity and vorticity of the fluid in two regions: a circular region encompassing the flow around the whole embryo (Fig. S7B), and a rectangular region isolating the flow immediately above the mouth of the embryo (Fig. S7C). We found that in control embryos, the ratio of the velocity of flow towards the mouth compared to the velocity surrounding the entire embryo is 0.23 on average (Fig. 7E). Similarly, the average ratio of the vorticity near the mouth to that surrounding the entire embryo is 0.31 (Fig. 7F).

In comparison, we observed two distinct flow patterns surrounding AZ-treated embryos depending on their morphology, which we classified as AZ I or II (Fig. 7B-C). AZ-treated embryos with the stereotypic long but abnormally wide aboral arms (AZ I) produced only two vortices (Fig. 7B, D). Embryos that had a more severe phenotype (AZ II) lacked aboral arms but still generated about six vortices (Fig. 7C, D). Compared to control, AZ-treated embryos had slightly higher overall velocity of fluid flow (Fig. 7B2, E); however, the velocity of fluid flow near the mouth was lower (Fig. 7E). Interestingly, the ratio of fluid velocity near the mouth versus the fluid velocity surrounding the whole embryo was significantly lower in both classes of AZ-treated embryos (Fig. 7E3).

Although there was no significant difference in overall vorticity in AZ-treated embryos, the average vorticity near the mouth was significantly reduced with AZ (Fig. 7F1-2). Like the flow velocity, the ratio of vorticity near the mouth relative to the entire embryo was significantly lower in AZ-treated embryos (p < 0.03, *t*-test) (Fig. 7F3). Taken together, these data suggest that fluid movements near the mouth are substantially reduced in AZ-treated embryos.

The PIV studies suggested that AZ-exposed embryos have compromised feeding activity. To test that prediction, we performed feeding assays by exposing control and AZ-treated larvae to *Rhodomonas* algae that naturally exhibit red fluorescence.

Because AZ itself also has red fluorescence that labels the skeleton, we included xylenol orange (XO) with the controls to produce a comparable effect. We quantified gut-specific fluorescence (Fig. 7G-I), and the results show that AZ-exposed embryos consume significantly less algae than controls, with a difference greater than 3-fold (Fig. 7J). These findings agree with the predictions from the PIV experiments, corroborating their reliability. Together, these findings demonstrate that abnormalities in larval morphology and neuronal connectivity induced by AZ treatment causes changes in ciliary-directed fluid flow that have functional consequences for larval feeding behaviors.

### AZ-mediated skeletal patterning can be phenocopied by catalase inhibition and transient H_2_O_2_ treatment

AZ has previously been reported to inhibit catalase activity (Hu et al., 2019).

Catalase enzyme catalyzes the breakdown of reactive oxygen species (ROS) such as hydrogen peroxide (H_2_O_2_) into water and oxygen. ROS are a normal byproduct of many metabolic pathways and have roles as signaling molecules; however, excessive amounts of ROS negatively affect macromolecule integrity and cell signaling, and the resulting oxidative stress contributes to cancers, autoimmune, and cardiovascular disorders, along with neurodegenerative diseases (Pham-Huy et al., 2008; Auten and Davis, 2009; Nandi et al., 2019). ROS imbalance also has negative impacts on development in vertebrate species (Auten and Davis, 2009). To explore this as a potential mechanism for the AZ-mediated perturbations in sea urchin embryos, we first checked the temporal expression dynamics of Lv-Catalase within our developmental transcriptome (Hogan et al., 2020). Catalase gene expression increases gradually during blastula stage and is highest during mid-gastrula and pluteus stages (Fig. 8A).

**Figure 8.**
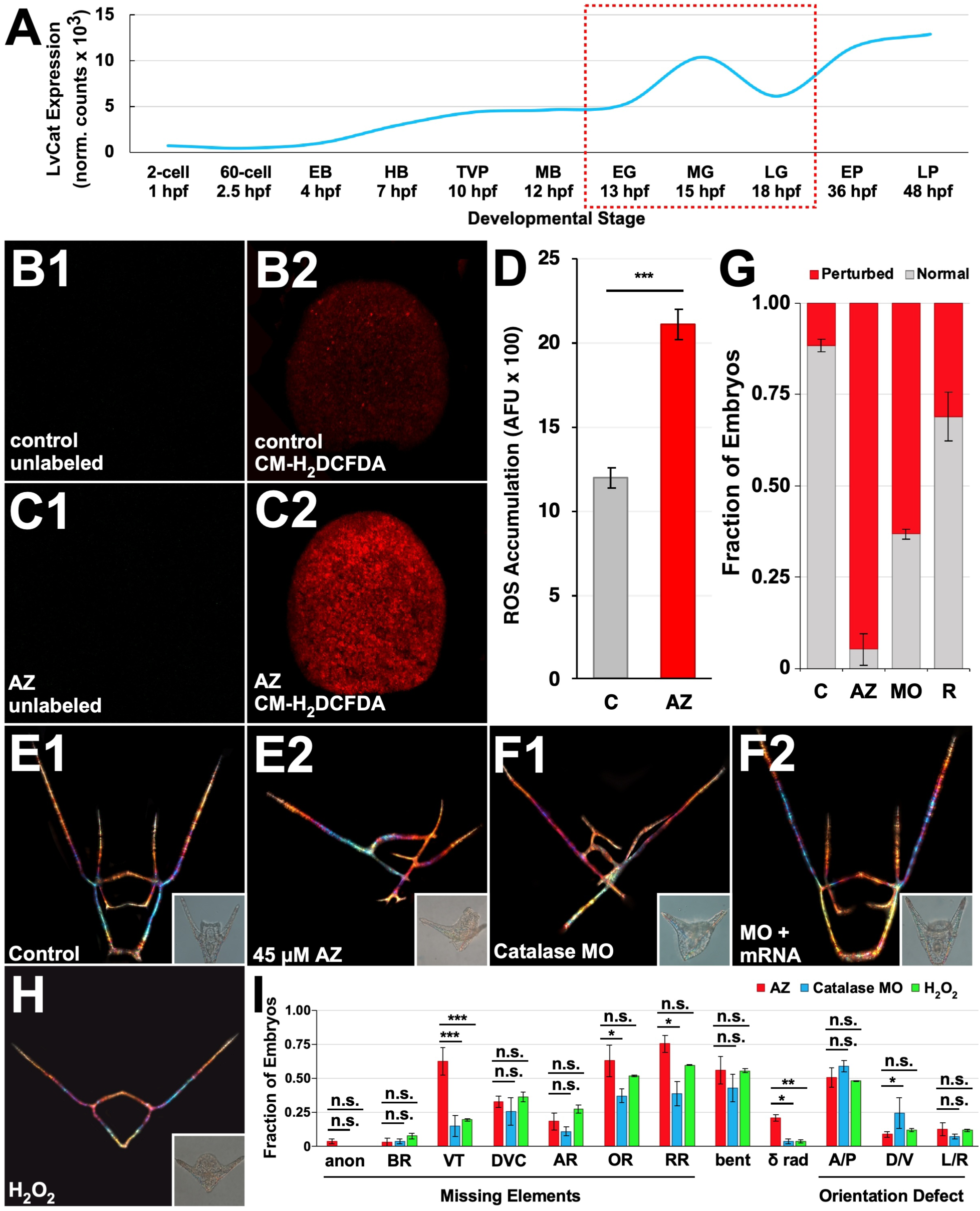
AZ-mediated skeletal patterning defects are phenocopied by catalase inhibition and H_2_O_2_ treatment. **A.** The level of Lv-Catalase expression is shown at the indicated stages as normalized reads from temporal transcriptomics data (Hogan et al., 2020). The red box indicates the window of effect of AZ. EB early blastula; HB hatched blastula; TVP thickened vegetal plate; MB mesenchyme blastula; EG early gastrula; MG mid gastrula; LG late gastrula; EP early pluteus; LP late pluteus. **B-C.** Representative control (B) and AZ-treated (C) embryos unlabeled (1) or stained with 20 μM CM- H_2_DCFDA (2) are shown at 12 hpf. **D.** Quantification of relative ROS levels is shown as average artificial fluorescence units (AFU) ± S.E.M. in control and AZ-treated embryos labeled with CM-H_2_DCFDA; n ≥ 113; *** p < 0.0005. **E-F.** Representative skeletal birefringence images of control (E1), AZ-treated (E2), catalase MO-injected (F1), and catalase MO and mRNA co-injected (F2) embryos are shown at 48 hpf. Insets show morphologies of the corresponding embryo visualized via DIC illumination. **G.** Fraction of embryos with perturbed skeletal patterning in control (C), AZ-treated (AZ), catalase MO-injected (MO), or catalase MO and mRNA co-injected (R) embryos is shown as average percentage ± S.E.M; n ≥ 57. **H.** Representative skeletal birefringence and morphology (DIC, inset) images at 48 hpf of an embryo treated with 50 μM H_2_O_2_ from 8- 10 hpf. **I.** The fractions of AZ-treated (red), catalase MO-injected (cyan), or H_2_O_2_-treated (green) embryos with the indicated skeletal patterning defects are displayed as the average percentage ± S.E.M; n ≥ 25; * p < 0.05; ** p < 0.005; *** p < 0.0005; n.s. not significant (*t*-test). Skeletal elements are abbreviated as in Fig. 1.

This peak of catalase expression during gastrulation matches the window during which AZ exposure is most effective (Fig. 1I-J). We next visualized ROS in *L. variegatus* embryos at 12 hpf with the fluorescent sensor CM-H_2_DCFDA and found that control embryos have minimal ROS while AZ treatment significantly elevated ROS throughout the embryo (Fig. 8B-D). This suggests that AZ inhibits catalase activity in sea urchin embryos, resulting in ROS accumulation.

To directly test if AZ-mediated inhibition of catalase contributes to the skeletal defects observed in AZ-treated embryos, we employed a morpholino-substituted antisense oligo (MO) designed to sterically block translation of *Lv*-catalase mRNA and thereby decrease production of Catalase enzyme. We found that skeletal patterns produced by *Lv*-catalase MO-injected zygotes exhibited the dramatic A-P rotational defects, bent skeletal rods, and missing skeletal elements that are characteristic of AZ-treated embryos (Fig. 8E-F, I). These defects were rescued by co-injection of catalase mRNA, confirming that MO specificity (Fig. 8F-G).

These results imply that excess levels of H_2_O_2_ contribute to the skeletal defects elicited by AZ. To test that hypothesis, we investigated whether H_2_O_2_ treatment phenocopies AZ treatment. Long-term H_2_O_2_ exposure was lethal (data not shown); so, we tested the effects of transient H_2_O_2_ exposure prior to gastrulation, immediately preceding the temporal window that matches peak sensitivity to AZ (Fig. S8A). We compared a range of concentrations of H_2_O_2_ during several two-hour windows and scored the relative abundance of normal, AZ-like, toxic, or otherwise perturbed embryos. The highest tested concentration, 60 μM, produced mostly toxic phenotypes during all treatment windows (Fig. S8E-F), while the other tested concentrations exhibited results that depended on the timing of treatment. Younger embryos exhibit greater sensitivity to ROS-mediated damage, which is disastrous before 10 hpf, then better tolerated after 10 hpf (Fig. S8B-F). This is kinetically consistent with the temporal window of sensitivity to AZ that we defined above.

We selected the 50 μM H_2_O_2_ dose with an 8-10 hpf treatment window for more in-depth scoring, since this treatment produced the largest percentage of AZ-like embryos and relatively low toxicity. Embryos treated in this manner had the dramatic A-P rotational defects, bent skeletal rods, and missing skeletal elements characteristic of AZ-treated embryos (Fig. 8H-I). We found no significant differences between prevalence of any of the scored phenotypes in catalase MO-injected embryos and H_2_O_2_-treated embryos (p > 0.05, *t*-tests). The loss of VTs and partial radialization present in some

AZ-treated embryos were not phenocopied by either catalase knockdown or transient H_2_O_2_ exposure (Fig. 8I), suggesting that these defects result from a different mechanism. Together, these data show that the majority of the effects of AZ treatment on skeleton formation can be accounted for by the inhibition of catalase and the concomitant increase in ROS.

### Model

The results presented in this study suggest a model in which AZ inhibits catalase, resulting in ROS accumulation and likely perturbing a wide range of processes, including gene expression, via the indiscriminate oxidizing actions of ROS. Our results indicate that PMC organization and skeletal patterning are particularly sensitive to these effects, while DV specification is impacted mildly and with lower penetrance.

Development overall is also substantially delayed by AZ exposure. We observed mild abnormalities in serotonergic neural process formation or extension with AZ treatment; together, these changes appear to have disrupted the integrity of cilia entrainment into uniform motion, based on the disruptions we observed in the fluid flow patterns around the larvae and their impaired feeding behavior. Our results also indicate that some of the effects of AZ, specifically the formation of the ventral primary skeletal elements and supernumerary triradiates, occur independently of oxidative dysregulation; however, the majority of AZ-mediated skeletal patterning defects can be accounted for by catalase inhibition and ROS accumulation.

## Discussion

Our findings reveal teratogenic effects of AZ treatment. Previous work has reported deformities that arise from AZ exposure in several species, including sea urchin larvae (Hoyte, 1960; Adkins, 1965; Rubin and Bisk, 1969; Lampertsdörfer et al., 1991; Descoteaux et al., 2023); however, our study is the first to characterize specific skeletal patterning and developmental defects resulting from AZ treatment. We show that AZ treatment dramatically perturbs skeletal patterning in sea urchin larvae, resulting in abnormal rotation and branching of the skeletal triradiates, bent skeletal rods, and missing skeletal elements. We also establish that these teratogenic effects are largely due to H_2_O_2_ accumulation following catalase inhibition by AZ, since transient H_2_O_2_ treatment prior to gastrulation and knockdown of catalase via MO injection both phenocopy most AZ-mediated skeletal patterning defects. Thus, we conclude that ROS are sufficient to disrupt skeletal patterning. Although the effects of ROS on early sea urchin development have previously been explored (Agca et al., 2009; Coffman et al., 2009; Chang et al., 2017), our study is the first to define a role for ROS later during skeletogenesis. This is also the first study to directly test the effect of AZ treatment on the accumulation of H_2_O_2_ during larval development and its consequences in the context of skeletal patterning.

The abnormal bending of the skeletal elements observed herein is a particularly interesting and unusual phenotype. This could simply be due to abnormal patterns of PMC migration caused by loss of PMC identity and/or ectodermal patterning cues, which we discuss below. Another possibility is that the ROS itself is acting as a cue that the PMCs respond to: previous studies in other systems have found that H_2_O_2_ can influence cell migration by acting as a chemoattractant, such as during wound healing (Klyubin et al., 1996; Niethammer et al., 2009; Hurd et al., 2012). Global AZ-induced ROS elevation might contradict membrane protein and extracellular matrix-based cues, thereby perturbing the direction of PMC migration and resulting in the seemingly random direction of skeletal elongation observed with each of these three perturbations. Yet another possibility is that ROS are directly affecting the crystallization or crystal structure of the larval skeleton by oxidizing various components of the skeletal matrix, with spicule matrix proteins being the most obvious candidates (Killian and Wilt, 1996; Zhu et al., 2001; Killian and Wilt, 2017). Structural studies will be necessary to elucidate the direct effect of AZ and/or ROS on the structure and composition of the skeletal matrix.

One possible way that ROS could mediate the effects of AZ on skeletal patterning and PMC migration is through changes to PMC identity and/or ectodermal patterning cues. ROS are well known to affect gene expression. Here, we find that AZ in sea urchin larvae has widespread effects on gene expression across a variety of gene categories and cellular compartments. Specifically, we find that AZ treatment results in abnormal expression level and/or location of several PMC subset genes, including *jun*, *pks2*, *sm50*, and *nk7*. The global activation of *jun* could reflect a temporal delay, since *jun* is initially globally expressed but is later spatially restricted to the PMCs (Howard-Ashby et al., 2006; Russo et al., 2014; Rodríguez-Sastre et al., 2023). Another possibility is that *jun* is activated globally as a stress response, since ROS, including H_2_O_2_, activate c-Jun in several vertebrate systems (Inanami et al., 1999; Chen et al., 2001; Kaneto et al., 2002; Ruffels et al., 2004; Touyz et al., 2004; Zhang et al., 2007). The elevation, but not spatial expansion, that we observe in expression of *pks2* may similarly reflect a temporal delay or loss of PMC identity, since *pks2* is initially expressed throughout all PMCs before being restricted to specific populations (Zuch and Bradham, 2019). Alternatively, perturbed *pks2* expression may also be a stress response, since other polyketide synthases are activated in response to oxidative stress in bacterial species (Jeanjean et al., 2008; Zhang et al., 2013). The connections between AZ and/or ROS and *sm50* and *nk7* has not been explored and are therefore novel observations.

Interestingly, we did not observe significant changes to the expression of known ectodermal patterning genes *vegf*, *univin*, or *wnt5* (Duloquin et al., 2007; Adomako-Ankomah and Ettensohn, 2013; McIntyre et al., 2013; Piacentino et al., 2015). This was surprising, since previous studies in other systems have reported that VEGF and Wnt5 are activated by ROS (Chua et al., 1998; Cho et al., 2001; Funato et al., 2006; Wang et al., 2016). However, it seems apparent that these pathways are simply not involved here, and that impacts on other ectodermal patterning cues more likely mediate the effects of AZ on the mis-patterning and loss of identity of the PMCs in sea urchin larvae. It is also possible that AZ directly impacts the PMCs and perturbs gene expression within them; that could further imply that PMCs are particularly sensitive or vulnerable to ROS elevation.

Although gross ectodermal specification is not affected in most AZ-treated embryos, we noted a profound effect of AZ treatment on the connectivity of serotonergic neurons. This population of neurons is thought to be the central nervous system of the sea urchin larva and to regulate ciliary beating (Bisgrove and Burke, 1986; Yaguchi et al., 2006; Angerer et al., 2011). The balance of ROS plays a critical role in neuronal development and maintenance. Although previous studies in vertebrate systems have reported that ROS such as H_2_O_2_ are required for axonal growth and branching (Munnamalai and Suter, 2009; Gauron et al., 2016), oxidative stress from excessive ROS can contribute to axon degeneration (Press and Milbrandt, 2008; Fukui et al., 2012) and to neurodegenerative diseases like Alzheimer’s and Parkinson’s (Behl et al., 1994; Matsuoka et al., 2001; Dias-Santagata et al., 2007; Shen et al., 2008; Cristóvão et al., 2012; Pukaß and Richter-Landsberg, 2014). Elevated ROS therefore may have similar neurodegenerative effects in sea urchin larvae, which would have consequences for ciliary-directed swimming behaviors.

Our PIV studies with confined larvae show that AZ exposure dramatically perturbs cilia-mediated fluid flow patterns around the embryo. In both control and AZ-treated larvae, the observed flow patterns along the body of the larvae appear to contribute to swimming. In control larvae, flows directed towards the mouth from vortices that form between the long arms facilitate feeding. Interestingly, AZ-exposed embryos with wide arms exhibit distinct patterns that lack flow at the mouth, while AZ- exposed embryos without arms exhibit weak fluid flows overall and near the mouth.

These changes in fluid flow patterns correlate with very limited feeding by AZ-exposed embryos, indicating that the flow patterns in control plutei indeed direct food to the larval mouth. The difference in both flow patterns and feeding efficiency between the controls and the wide-arm AZ-exposed larvae suggests that the normal form of the pluteus skeleton, with the long arms at an angle of approximately 45°, optimizes feeding by directing fluid flow to the mouth. This finding provides evidence to support the idea that the pluteus larval skeletal structure is an example of convergent evolution because it is an ideal structure for feeding, as has been previously speculated (Strathmann, 2006).

Although many of the AZ-mediated defects can be explained by H_2_O_2_ elevation due to catalase inhibition, some defects notably cannot; in particular, neither the loss of the VTs nor the presence of supernumerary triradiates is phenocopied by catalase knockdown or H_2_O_2_ treatment. This suggests that these defects are due to a different mechanism. The specific lack of phenocopy of VT defects is particularly interesting, since loss of other skeletal elements in consistent across all three perturbations. We have shown that AZ treatment results in changes at the genetic level, but it may also exert changes at the biophysical level. AZ is a calcium-binding fluorochrome, meaning that its direct interaction with calcium could be affecting the ion’s bioavailability for incorporation into the biomineral, or affecting its coordination in the crystal structure. No studies to date have explored the variation in crystal structure or biomineral composition of the different skeletal elements; however, given the differences in shape and curvature of the skeletal elements, it is possible that differences in the biomineral exist; that prediction is further supported by spatially differential expression of spicule matrix protein-encoding genes such as SM30 and SM50 among the PMCs. Perhaps the VTs are more sensitive to calcium bioavailability and thus are the first to be lost when calcium is sequestered.

Our study is among the earliest to connect AZ exposure to development defects and their underlying mechanisms and reveals that AZ is a teratogen. Use of AZ, particularly in textile or other large-scale commercial applications therefore presents a serious environmental concern. Because AZ is highly water-soluble, it can readily pollute aquatic systems if wastewater is not properly treated. Currently, no effective large-scale methods for removing AZ from wastewater exist. Although decolorization is an active area of research, the dye remains difficult to remove and its use and disposal are poorly regulated (Devi et al., 2009; Mahmoodi and Arami, 2009; Fan et al., 2012; Ghaedi et al., 2012; Roosta et al., 2014; Adnan et al., 2017; Pourebrahim et al., 2017; Zhang et al., 2019; Jian et al., 2021; Zhang et al., 2021; Vedula and Yadav, 2023). Our study shows how the presence of AZ in sea water could have adverse effects on the development of aquatic species, emphasizing the need for regulation of alizarin disposal and advancement of decolorization techniques.

## Materials and Methods

### Animals and embryo cultures

Adult *L. variegatus* sea urchins were obtained either from Laura Salter (Davis, NC) or Reeftopia (Miami, FL) and spawned via intraperitoneal injection of 0.5 M potassium chloride to collect gametes. Embryos were cultured at 23 °C in artificial sea water (ASW).

### Mineralization markers and polychrome labeling

Alizarin red S (AZ, Sigma #A5533), xylenol orange tetrasodium salt (XO, Sigma #398187), calcein green (CG, ThermoFisher #C481), and calcein blue (CB, Sigma #M1255) stocks were prepared separately by dissolving in distilled water.

Concentrations used were 45 μM AZ, 30 μM XO, 30 μM CG, and/or 45 μM CB, unless otherwise indicated. Polychrome labeling experiments with 24 hpf color switches were performed as previously described (Descoteaux et al., 2023), with the exception of simultaneous incubation in XO for the duration of the experiment in controls to account for the simultaneous presence of two fluorochromes in AZ-treated embryos.

### Immunostains

Embryos were fixed and immunostained as previously described (Bradham et al., 2009). PMC-specific primary antibody 6a9 (1:30) was a gift from Charles Ettensohn (Carnegie Mellon University). Ciliary band-specific primary antibody 295 (undiluted) was a gift from David McClay (Duke University). Anti-serotonin (Sigma) was used to label serotonergic neurons. Goat anti-mouse Alexa 488 (ThermoFisher, 1:500) was used as secondary antibody. Hoechst 33258 (Sigma) was used to label nuclei (1:1000).

### Hybridization Chain Reaction Fluorescent in situ Hybridization (HCR FISH)

Embryos were fixed at the indicated time points in 4% paraformaldehyde and HCR FISH was performed in 1.5 mL microcentrifuge tubes following the published protocol for sea urchin embryos (https://files.molecularinstruments.com/MI-Protocol-RNAFISH-SeaUrchin-Rev9.pdf). Probe sets were designed from open reading frames of the indicated genes by Molecular Instruments, Inc (Los Angeles, CA). Probes were used at the recommended 8 nM concentration with the following exceptions: VEGFR, Univin, Wnt5 (16 nM), and Jun (32 nM). Hairpins tagged with Alexa 488, Alexa 546, or Alexa 647 were used to amplify and fluorescently label the transcripts of the genes of interest.

### Transcriptomics

Total RNA was prepared from 10,000 embryos as previously described (Rodríguez-Sastre et al., 2023). In each case, cultures were divided into control and AZ- treatments; paired samples were collected simultaneously. Bulk library preparation and sequencing from biological triplicates was performed using DNB (Innomics, Inc.) and Illumina sequencing, respectively. In some cases, total RNA samples were replaced due to library failure. Quality control of the resulting data was performed using FastQC v0.12.1-0 (bioinformatics.babraham.ac.uk/projects/fastqc/). The reads were then aligned to the Lvar 3.0 genome using STAR 2.7.1 and the read counts were calculated using Verse 0.1.5 (Davidson et al., 2020; Dobin et al., 2013; Zhu et al., 2016). Raw fastq files and the processed count matrix are available at GEO (accession number GSE270804). Multiple gene models mapping to a single gene annotation were collapsed to a single entry by summing read counts across models. Quantile normalization was performed using the preprocessCore package in R (github.com/bmbolstad/preprocessCore), then differential gene expression analysis was performed pairwise using the DESeq2 package in R with the size factors were explicitly set to 1 to prevent DESeq2 from performing a second normalization (Love et al., 2014). Principal component analysis (PCA) was performed using the PCAtools package (github.com/kevinblighe/PCAtools) and GO Enrichment analysis was performed using the gseapy package in Python (Fang et al., 2023). Enriched and depleted GO terms were manually binned into broad categories (see Fig. 4B). GO enrichment results were semantically clustered using Revigo, then visualized and merged using cytoscape (Kucera et al., 2016; Supek et al., 2011). The corresponding code for these analyses can be found at https://github.com/BradhamLab/AZ-paper-analysis-code.

### ROS detection

H_2_O_2_ detection in live embryos with CM-H_2_DCFDA (ThermoFisher #C6827) was performed as previously described (Coffman et al., 2009). Embryos were knocked down with 2X ASW at 12 hpf and labeled with 10 μM CM-H_2_DCFDA for 20 minutes before confocal imaging.

### Microscopy

Skeletal birefringence and morphology differential interference contrast (DIC) images of larvae were collected using a Zeiss Axioplan upright microscope at 200x magnification. For skeletal images, multiple focal planes were imaged using plane-polarized light and montaged in ImageJ. Out-of-focus light was manually removed so that the entire skeleton was in focus. Confocal z-stacks were collected using a Nikon C2Si laser-scanning confocal microscope and projected maximally in 2-D or 3-D using ImageJ. Embryos were imaged with identical acquisition settings within each experiment to ensure their comparability.

### Flow field imaging and analysis

To quantify the fluid flows and their patterns around control and AZ-treated sea urchin larvae, we performed live imaging experiments under squeeze-confinement conditions. Live embryos were mounted on slides using 50 µm double-stick tape spacers (Nitto Inc.) in ∼ 50 µL of ASW. 1 µm polystyrene microspheres (Polysciences, cat. no. 07310-15) in a second droplet (∼ 10 µl) were added and gently mixed for a final dilution of ∼ 1:100. After adding a coverslip, the sea urchin larvae were gently trapped under 50 µm squeeze-confinement conditions.

To quantify the movement of the microspheres (induced by the larval ciliary beating), we carried out live darkfield imaging with an upright microscope (Zeiss Axio Imager M2) using a 10X objective that captures a 1.5 mm x 1.5 mm field of view. We captured time-lapse images at 10 frames per second for 30 seconds using a high-speed camera (Hamamatsu ORCA-Fusion Gen-III sCMOS). To quantify the larval fluid flows, time-lapse images were first post-processed using PIVlab in MATLAB (Thielicke and Stamhuis, 2014). A mask was drawn over the larva in PIVlab to avoid any flow quantification artifacts inside the body due to noise. We next removed background noise using high-pass filters, then calculated the time-varying velocity vector fields. The velocity vector fields were post-processed to smoothen the fields and spatially interpolate vectors when necessary. For smooth visualizations of fluid flows and for quantification of the results, we calculated the averaged mean of the parameter of interest over 30 frames, corresponding temporally to 3 seconds. To eliminate background, whole embryo quantification was limited to a 200-micron diameter circular region centered on the larva (Fig. S7B). Flow near the mouth was quantified in a boxed region encompassing the mouth (Fig. S7C). The flow parameters quantified in Fig. 7E-F were calculated by summing the values in the appropriate region.

### Feeding experiments

Following optimization, embryos were incubated with 20,000 cells/mL of *Rhodomonas* algae that naturally exhibit red fluorescence at 24 hpf (early pluteus stage), then washed with ASW three times before imaging at 48 hpf. To match AZ-mediated red fluorescent labeling of the skeleton, the skeletons in control embryos were comparably labeled with xylenol orange (XO) (Descoteaux et al., 2023). Relative fluorescence intensities were quantified in FIJI as the mean gray value per unit area; net fluorescence values were averaged across three biological replicates.

### Scoring and statistics

At least three biologically independent replicates were used for each experiment unless otherwise noted. Skeletal scoring was performed on montaged skeletal images using an in-house scoring rubric (Piacentino et al., 2016a; Thomas et al., 2023).

Maximally projected confocal z-stacks of HCR FISH signals were used to quantify gene expression. Regions of interest (ROIs) were drawn around the entire embryo in ImageJ, and the mean fluorescence value per unit area was measured from each ROI as previously described (Thomas et al., 2023). Statistical significance was analyzed via paired Student’s *t*-tests using two-tailed heteroscedastic settings in Microsoft Excel. P-values less than 0.05 were considered significant.

## Author Contributions

This study was conceived and designed by CAB and AED. The experiments were executed by AED, MR, BDS, and DA; data analyses were performed by AED, MR, MG, BDS, VNP, and CAB; The manuscript was written by AED, MR and CAB, and edited by all co-authors.

## Acknowledgments

We thank Professors Charles Ettensohn and David McClay for their gifts of antibodies, and Dr. Todd Blute for microscopy advice. This work was supported by NSF IOS 1656752 (CAB) and NIGMS 1R35GM152180 (CAB). A.E.D. was partially supported by the Boston University Biological Design Center’s Kilachand Fellowship. M.R. was partially supported by the Boston University Biological Design Center STEM Pathways program (DoD STEM FY20 Award HQ00342110008) and the Boston University Undergraduate Research Opportunities Program (UROP).

