## Supplementary material for "Alizarin red perturbs skeletal patterning and biomineralization via Catalase inhibition": Fig. S1-S8

Figures S1-S8 (this document)

Table S1. GO enrichment (excel file)

Movie S1
A live control larva at 48 hpf whose skeleton was labeled with xylenol orange, shown rotating through all body axes. (mp4 file)

Movie S2
A live AZ-treated larva at 48 hpf whose skeleton was labeled with calcein blue, shown rotating through all body axes. (mp4 file)

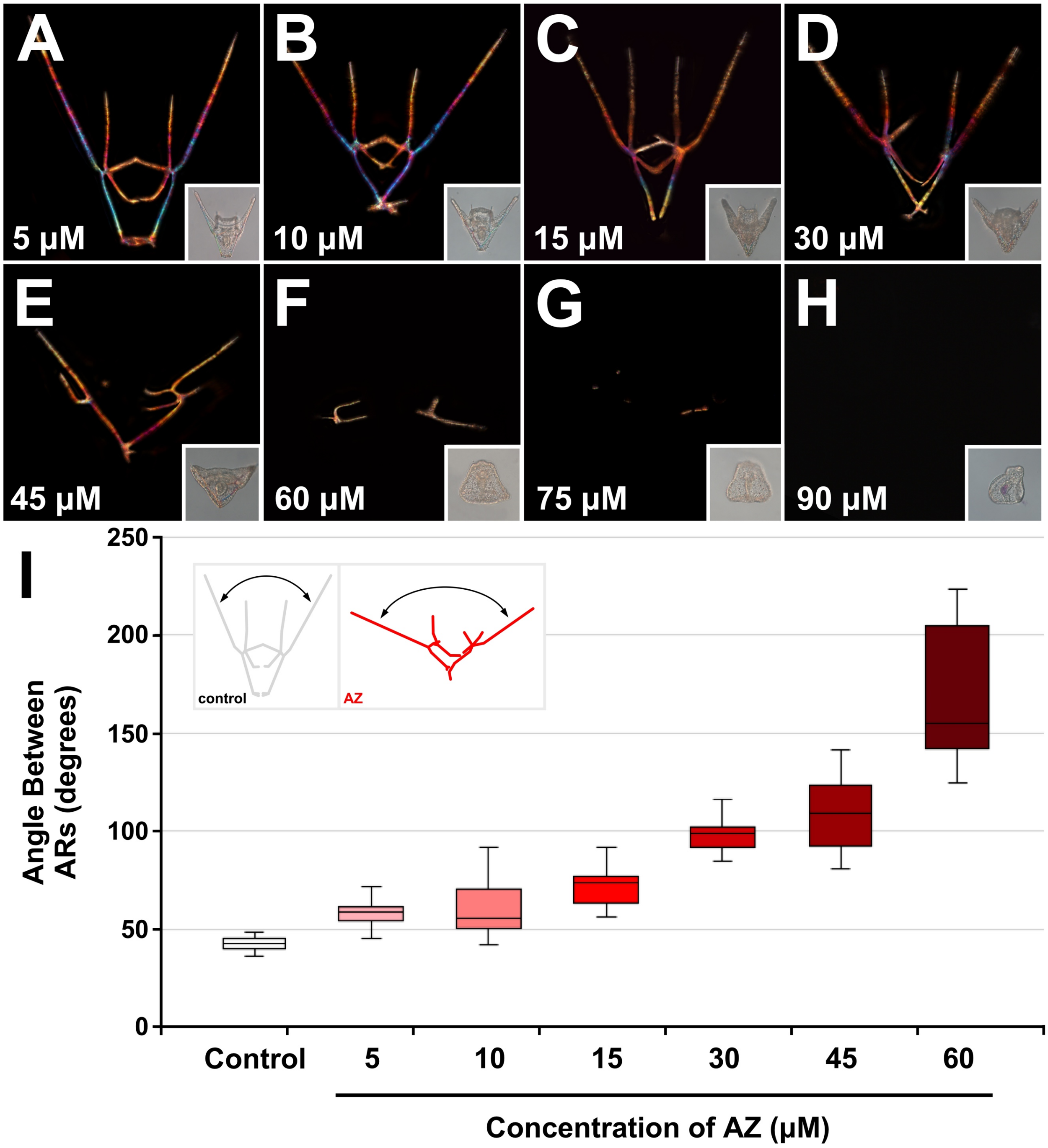


**Fig. S1. Defects caused by AZ treatment are dose-dependent. A-H.** Skeletal birefringence and morphology (insets) of exemplar embryos treated with the indicated concentrations (μM) of AZ from fertilization until imaging are shown at 48 hpf. **I.** The range of angle measurements between the aboral rods (ARs) are shown as box-and-whiskers plots, with the whiskers reflecting the 10$^{th}$ and 90$^{th}$ percentiles; $n \geq 14$. Inset shows angle being compared between control and AZ embryos.

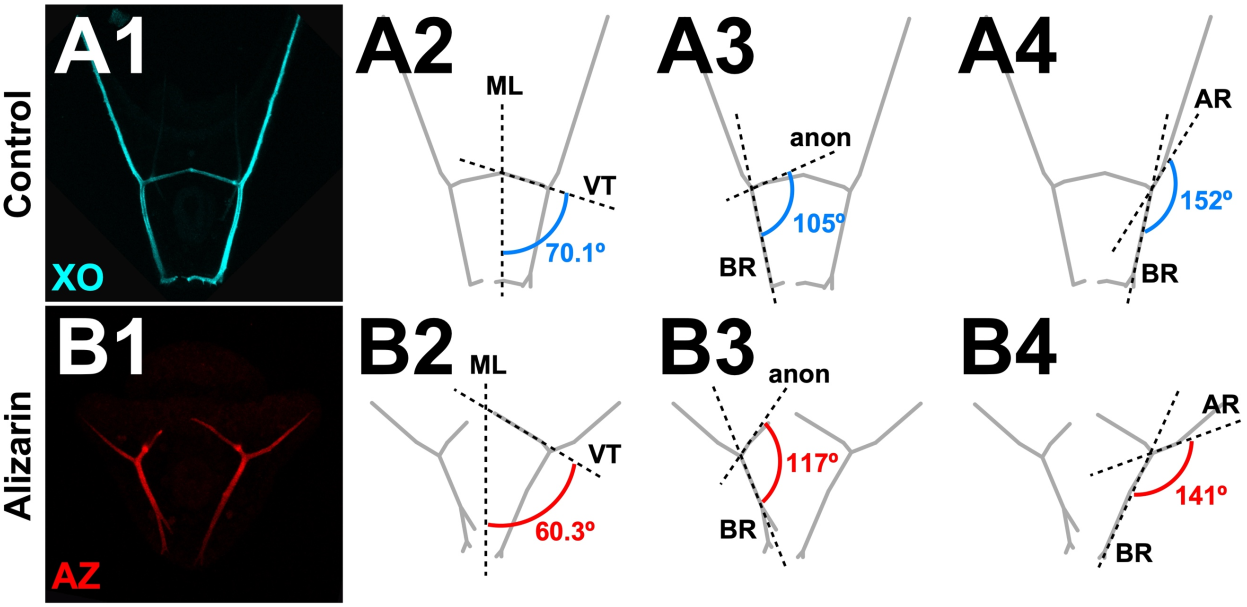


**Fig. S2. AZ treatments perturbs triradiate initiation, orientation, and branching. A-B.** A control XO-labeled (A) and an AZ-treated (B) embryo are shown (1). The average angles between the midline (ML) and ventral transverse rod (VT) (2), the body rod (BR) and anonymous rod (anon) (3), and the BR and aboral rod (AR) (4) are shown.

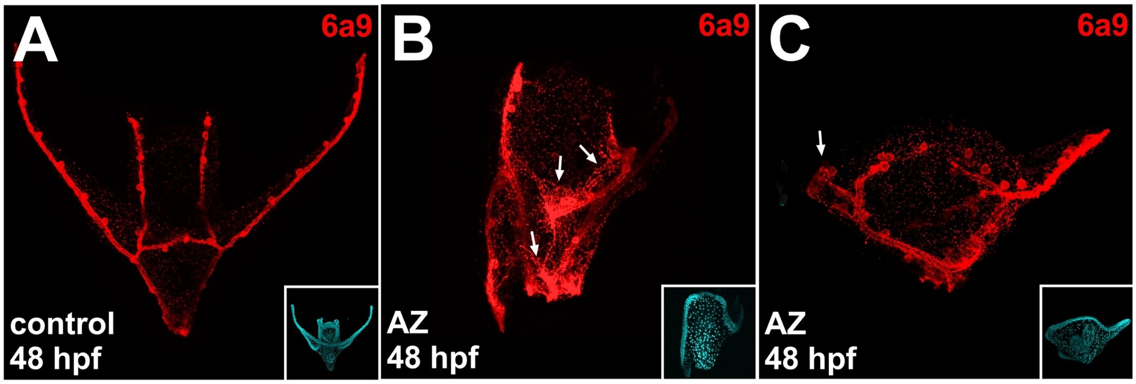


**Fig. S3. AZ treatment perturbs PMC migration and filopodial organization. A-C.** Control (A) and AZ-treated embryos (B-C) were immunolabeled at 48 hpf to visualize the PMCs. Insets show nuclei labeled with Hoechst in the corresponding embryo. Arrows in B indicate filopodial webbing. Arrows in C indicated a stalled plug of PMCs.

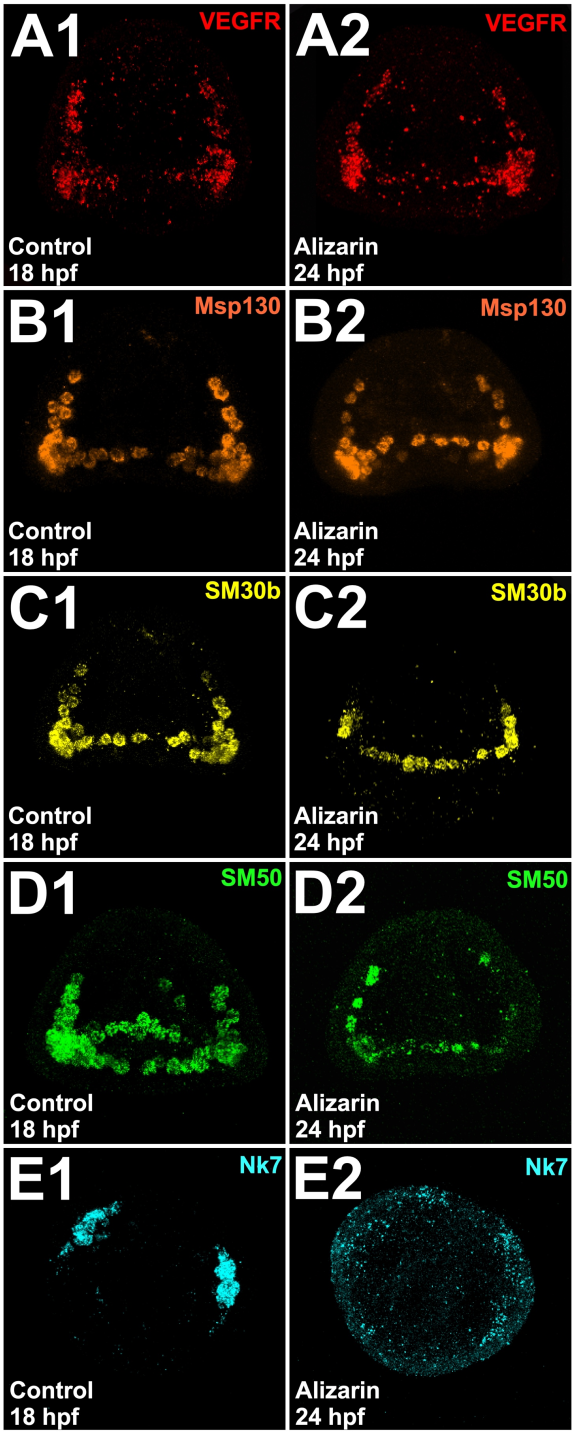


**Fig. S4. AZ treatment perturbs expression of some PMC subset genes. A-E.** Exemplar control (1) and AZ-treated embryos (2) that were subjected to HCR FISH at late gastrula stage for VEGFR (A), Msp130 (B), SM30b (C), SM50 (D), or Nk7 (E) are shown in *en face* (A-C), skewed (D), or vegetal (E) views.

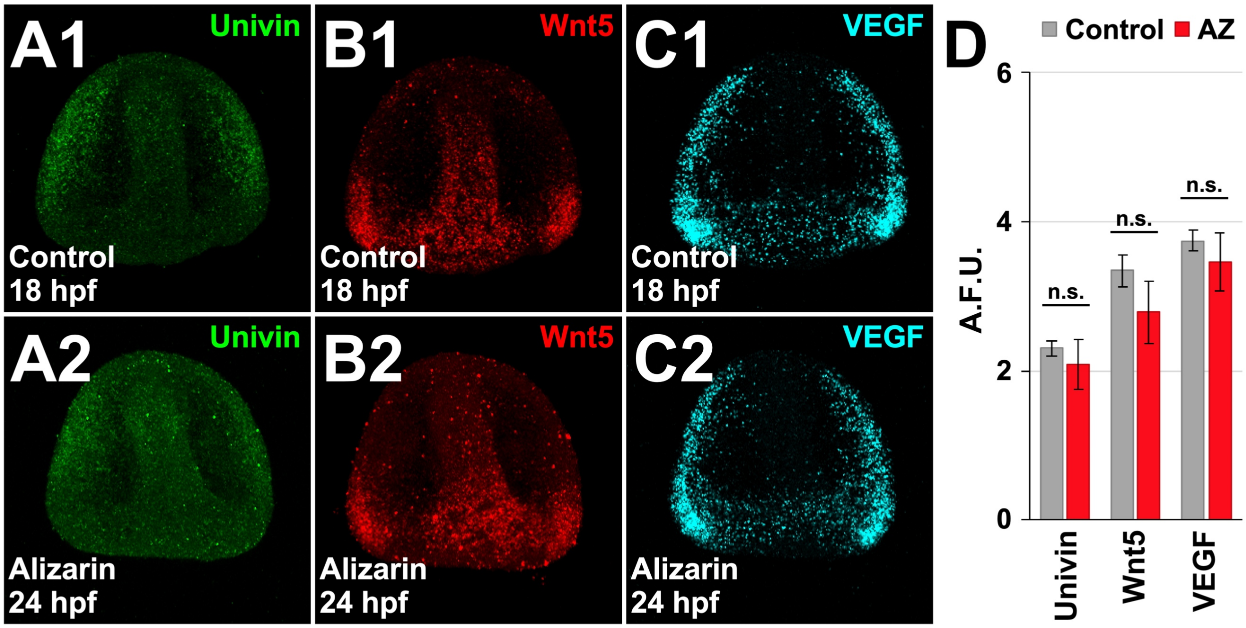


**Fig. S5. AZ treatment does not affect expression of known ectodermal patterning cues. A-C.** Exemplar control (1) and AZ-treated embryos (2) that were subjected to HCR FISH for Univin (A), Wnt5 (B), and VEGF (C) are shown at late gastrula stage. **D.** The normalized average expression per unit area of the indicated ectodermal genes is shown as the average artificial fluorescent unit (A.F.U.) ± S.E.M in control (grey) and AZ-treated embryos (red); n ≥ 19; n.s. not significant (*t*-test).

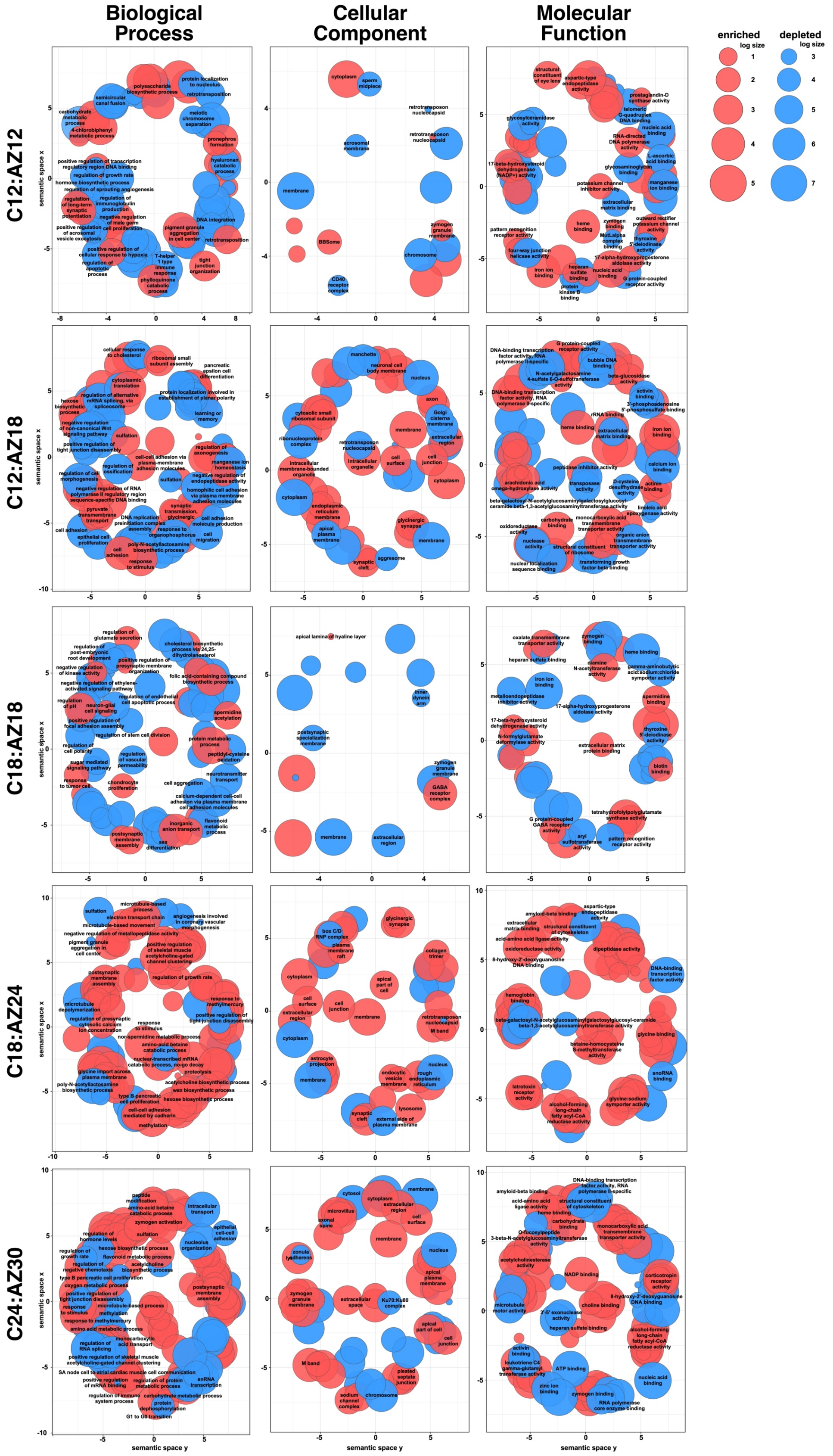
Biological Process
Cellular Component
Molecular Function
enriched
log size
depleted
log size
C12:AZ12
C12:AZ18
C18:AZ18
C18:AZ24
C24:AZ30
semantic space x
semantic space y

**Fig. S6. AZ treatment leads to broad changes in gene expression.** Semantic clustering of GO terms enriched and depleted by AZ relative to controls (C) is shown for two monochronic and three heterochronic comparisons at the indicated time points (hours post-fertilization).

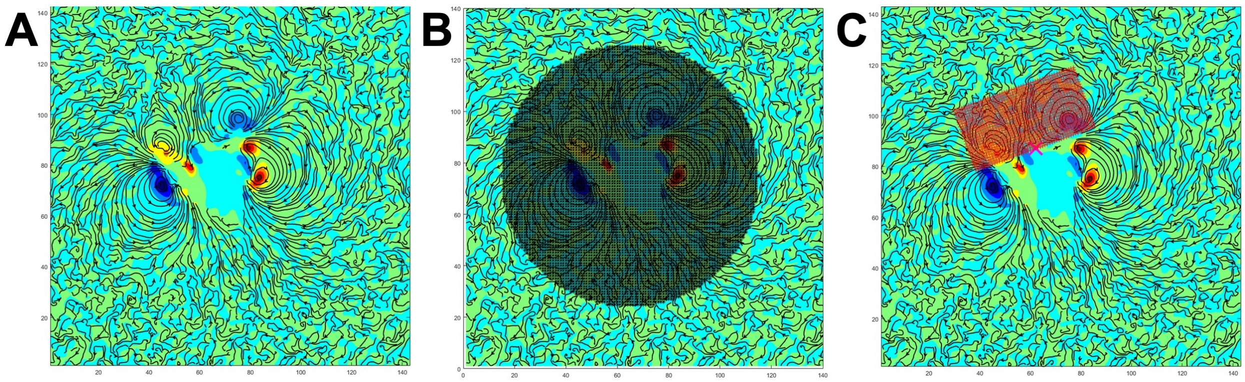

**Fig. S7. AZ treatment affects fluid vorticity around sea urchin larvae. A-C.** An exemplar flow field around a control embryo is shown with flow vectors (black) and shading depicting clockwise (red) or counterclockwise (blue) flow. The regions measured for overall fluid movements (B, grey circle) and local fluid movements near the mouth (C, red rectangle) are also shown.

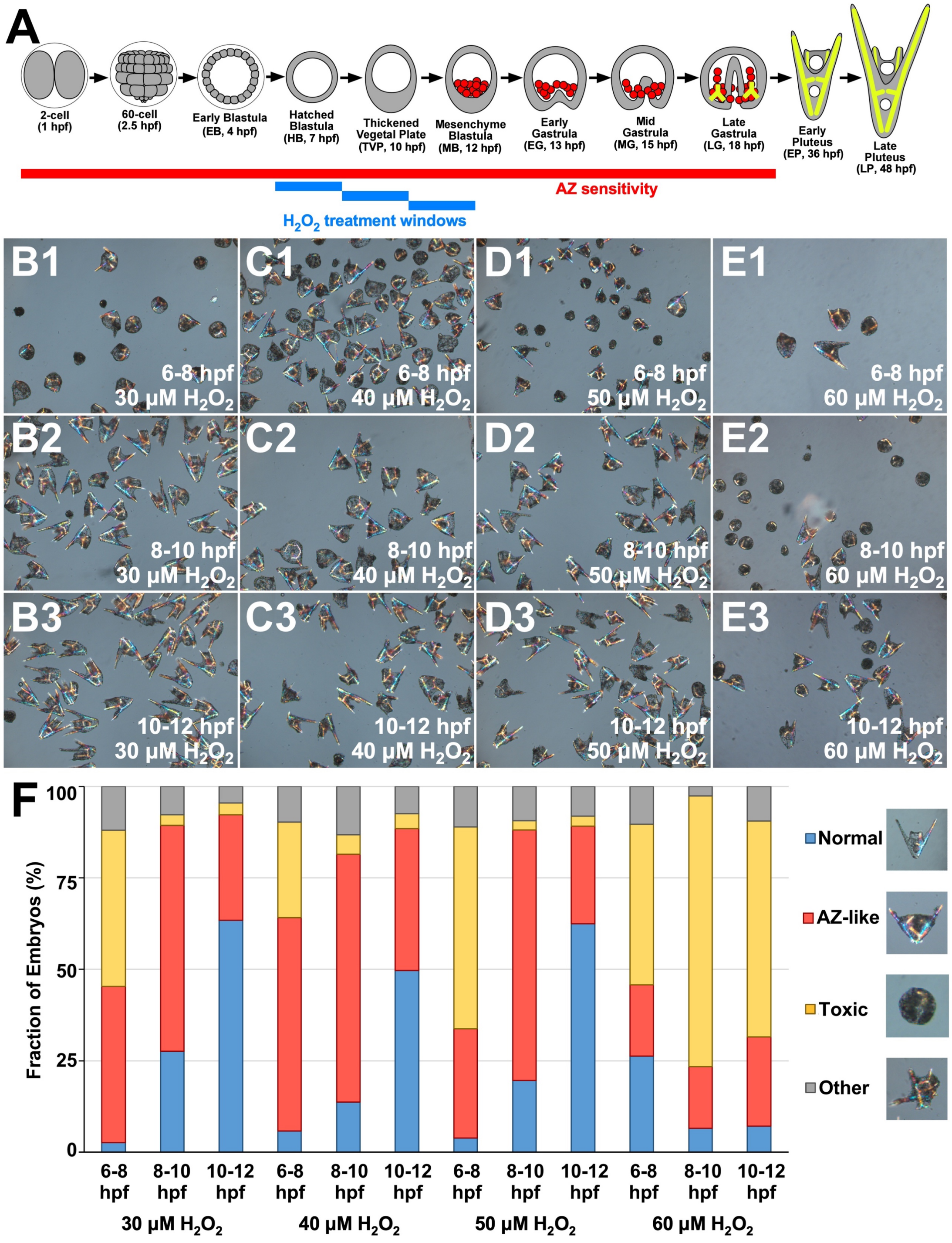


**Fig. S8. The effects of transient $H_2O_2$ treatment are dose-dependent and phenocopy AZ. A.** A timecourse showing the approximate windows of AZ sensitivity (red) and $H_2O_2$ treatment (blue) relative to larval development. The schematic (top) is adapted from (Hogan et al., 2020). **B-E.** Exemplar embryos were treated with 30 (B), 40 (C), 50 (D), or 60 µM (E) $H_2O_2$ from 6-8 hpf (1), 8-10 hpf (2), or 10-12 hpf (3), then imaged at 48 hpf. **F.** The fraction of embryos showing normal (green), AZ-like (yellow), toxic (blue), or otherwise abnormal development (grey) is shown as average percentage for each treatment condition; Normal: control-like pluteus skeleton; AZ-like: A-P rotational defects, bent skeletal rods, and/or missing anterior skeletal elements; Toxic: exogastrulation and/or round, compact morphology with little skeletal biomineralization or gastrulation and excessive mesenchymal accumulation; Other: Non-AZ-like phenotypes, such as D-V or L-R rotational defects and/or extra or duplicated skeletal elements; $n \geq 77$ per treatment interval.
