## Supplementary material for "Alizarin red perturbs skeletal patterning and biomineralization via Catalase inhibition": Table S1

| Term | Gene_set |
| --- | --- |
| GO:0005509 | calcium ion binding |
| GO:0005262 | calcium channel activity |
| GO:1901019 | regulation of calcium ion transmembrane transporter activity |
| GO:0090279 | regulation of calcium ion import |
| GO:1901021 | positive regulation of calcium ion transmembrane transporter activity |
| GO:0006874 | cellular calcium ion homeostasis |
| GO:0050839 | cell adhesion molecule binding |
| GO:1903037 | regulation of leukocyte cell-cell adhesion |
| GO:0005615 | extracellular space |
| GO:0050840 | extracellular matrix binding |
| GO:0005576 | extracellular region |
| GO:0031012 | extracellular matrix |
| GO:0005518 | collagen binding |
| GO:0030336 | negative regulation of cell migration |
| GO:0043034 | costamere |
| GO:0030054 | cell junction |
| GO:1903598 | positive regulation of gap junction assembly |
| GO:0044331 | cell-cell adhesion mediated by cadherin |
| GO:0045296 | cadherin binding |
| GO:0007155 | cell adhesion |
| GO:0005576 | extracellular region |
| GO:0005615 | extracellular space |
| GO:0062023 | collagen-containing extracellular matrix |
| GO:0005201 | extracellular matrix structural constituent |
| GO:0031012 | extracellular matrix |
| GO:0005581 | collagen trimer |
| GO:1904399 | heparan sulfate binding |
| GO:0005604 | basement membrane |
| GO:0030198 | extracellular matrix organization |
| GO:0010632 | regulation of epithelial cell migration |
| GO:0061580 | colon epithelial cell migration |
| GO:0031430 | M band |
| GO:0030286 | dynein complex |
| GO:0008569 | ATP-dependent microtubule motor activity, minus-end-directed |
| GO:0005856 | cytoskeleton |
| GO:0051015 | actin filament binding |
| GO:0051959 | dynein light intermediate chain binding |
| GO:0008307 | structural constituent of muscle |
| GO:0045505 | dynein intermediate chain binding |
| GO:0031672 | A band |

| | |
|---|---|
| GO:0030018 | Z disc |
| GO:0003779 | actin binding |
| GO:0030241 | skeletal muscle myosin thick filament assembly |
| GO:0071688 | striated muscle myosin thick filament assembly |
| GO:0005902 | microvillus |
| GO:0007018 | microtubule-based movement |
| GO:0005200 | structural constituent of cytoskeleton |
| GO:0051693 | actin filament capping |
| GO:0036309 | protein localization to M-band |
| GO:0030507 | spectrin binding |
| GO:0036156 | inner dynein arm |
| GO:0005859 | muscle myosin complex |
| GO:0005874 | microtubule |
| GO:0007275 | multicellular organism development |
| GO:0060216 | definitive hemopoiesis |
| GO:0030855 | epithelial cell differentiation |
| GO:0030326 | embryonic limb morphogenesis |
| GO:0009888 | tissue development |
| GO:0010831 | positive regulation of myotube differentiation |
| GO:0060122 | inner ear receptor cell stereocilium organization |
| GO:0061113 | pancreas morphogenesis |
| GO:0030858 | positive regulation of epithelial cell differentiation |
| GO:0097094 | craniofacial suture morphogenesis |
| GO:0001822 | kidney development |
| GO:0001525 | angiogenesis |
| GO:0000943 | retrotransposon nucleocapsid |
| GO:0043371 | negative regulation of CD4-positive, alpha-beta T cell differentiation |
| GO:0043378 | positive regulation of CD8-positive, alpha-beta T cell differentiation |
| GO:0000943 | retrotransposon nucleocapsid |
| GO:0009617 | response to bacterium |
| GO:0001618 | virus receptor activity |
| GO:0005044 | scavenger receptor activity |
| GO:0043152 | induction of bacterial agglutination |
| GO:0061844 | ntimicrobial humoral immune response mediated by antimicrobial peptid |
| GO:0038187 | pattern recognition receptor activity |
| GO:0035011 | melanotic encapsulation of foreign target |
| GO:0050829 | defense response to Gram-negative bacterium |
| GO:0050830 | defense response to Gram-positive bacterium |
| GO:0016021 | integral component of membrane |
| GO:0005886 | plasma membrane |
| GO:0009986 | cell surface |

| | |
|---|---|
| GO:0004180 | carboxypeptidase activity |
| GO:0016324 | apical plasma membrane |
| GO:0005887 | integral component of plasma membrane |
| GO:0042383 | sarcolemma |
| GO:0016020 | membrane |
| GO:0016323 | basolateral plasma membrane |
| GO:0030315 | T-tubule |
| GO:0007009 | plasma membrane organization |
| GO:0009897 | external side of plasma membrane |
| GO:0031526 | brush border membrane |
| GO:0033292 | T-tubule organization |
| GO:0045177 | apical part of cell |
| GO:0005903 | brush border |
| GO:0008270 | zinc ion binding |
| GO:0006491 | N-glycan processing |
| GO:0003828 | alpha-N-acetylneuraminate alpha-2,8-sialyltransferase activity |
| GO:0006486 | protein glycosylation |
| GO:0001574 | ganglioside biosynthetic process |
| GO:0010033 | response to organic substance |
| GO:0033829 | O-fucosylpeptide 3-beta-N-acetylglucosaminyltransferase activity |
| GO:0006470 | protein dephosphorylation |
| GO:0009311 | oligosaccharide metabolic process |
| GO:0008191 | metalloendopeptidase inhibitor activity |
| GO:0004190 | aspartic-type endopeptidase activity |
| GO:0017083 | 4-galactosyl-N-acetylglucosaminide 3-alpha-L-fucosyltransferase activity |
| GO:0004527 | exonuclease activity |
| GO:0008270 | zinc ion binding |
| GO:0004181 | metallocarboxypeptidase activity |
| GO:0042589 | zymogen granule membrane |
| GO:0035375 | zymogen binding |
| GO:0030246 | carbohydrate binding |
| GO:0050731 | positive regulation of peptidyl-tyrosine phosphorylation |
| GO:0070891 | lipoteichoic acid binding |
| GO:0050998 | nitric-oxide synthase binding |
| GO:0033265 | choline binding |
| GO:0000166 | nucleotide binding |
| GO:0032092 | positive regulation of protein binding |
| GO:0051117 | ATPase binding |
| GO:0005542 | folic acid binding |
| GO:0008289 | lipid binding |
| GO:0004553 | hydrolase activity, hydrolyzing O-glycosyl compounds |

| | |
|---|---|
| GO:0003943 | N-acetylgalactosamine-4-sulfatase activity |
| GO:0008484 | sulfuric ester hydrolase activity |
| GO:0004065 | arylsulfatase activity |
| GO:0006579 | amino-acid betaine catabolic process |
| GO:0051597 | response to methylmercury |
| GO:0016805 | dipeptidase activity |
| GO:0005975 | carbohydrate metabolic process |
| GO:0006508 | proteolysis |
| GO:0008194 | UDP-glycosyltransferase activity |
| GO:0008237 | metallopeptidase activity |
| GO:0008241 | peptidyl-dipeptidase activity |
| GO:0009268 | response to pH |
| GO:0008239 | dipeptidyl-peptidase activity |
| GO:0008238 | exopeptidase activity |
| GO:0047150 | betaine-homocysteine S-methyltransferase activity |
| GO:0043171 | peptide catabolic process |
| GO:0045329 | carnitine biosynthetic process |
| GO:0008292 | acetylcholine biosynthetic process |
| GO:0008233 | peptidase activity |
| GO:0004190 | aspartic-type endopeptidase activity |
| GO:0015020 | glucuronosyltransferase activity |
| GO:0004132 | dCMP deaminase activity |
| GO:0080019 | fatty-acyl-CoA reductase (alcohol-forming) activity |
| GO:0050129 | N-formylglutamate deformylase activity |
| GO:0102965 | alcohol-forming fatty acyl-CoA reductase activity |
| GO:0006196 | AMP catabolic process |
| GO:0051001 | negative regulation of nitric-oxide synthase activity |
| GO:0008206 | bile acid metabolic process |
| GO:0042632 | cholesterol homeostasis |
| GO:0150094 | amyloid-beta clearance by cellular catabolic process |
| GO:0003990 | acetylcholinesterase activity |
| GO:0004767 | sphingomyelin phosphodiesterase activity |
| GO:0050253 | retinyl-palmitate esterase activity |
| GO:0010025 | wax biosynthetic process |
| GO:0006766 | vitamin metabolic process |
| GO:0042426 | choline catabolic process |
| GO:0097242 | amyloid-beta clearance |
| GO:0070006 | metalloaminopeptidase activity |
| GO:0007584 | response to nutrient |
| GO:0052695 | cellular glucuronidation |
| GO:0005737 | cytoplasm |

| | |
|---|---|
| GO:0099699 | integral component of synaptic membrane |
| GO:0007399 | nervous system development |
| GO:0014704 | intercalated disc |
| GO:0045211 | postsynaptic membrane |
| GO:0031594 | neuromuscular junction |
| GO:0050896 | response to stimulus |
| GO:0042135 | neurotransmitter catabolic process |
| GO:0071340 | skeletal muscle acetylcholine-gated channel clustering |
| GO:0045989 | positive regulation of striated muscle contraction |
| GO:0002027 | regulation of heart rate |
| GO:0007605 | sensory perception of sound |
| GO:0098885 | modification of postsynaptic actin cytoskeleton |
| GO:0060048 | cardiac muscle contraction |
| GO:0045494 | photoreceptor cell maintenance |
| GO:0045202 | synapse |
| GO:0014819 | regulation of skeletal muscle contraction |
| GO:0043049 | otic placode formation |
| GO:0086070 | SA node cell to atrial cardiac muscle cell communication |
| GO:0086066 | atrial cardiac muscle cell to AV node cell communication |
| GO:0086036 | regulation of cardiac muscle cell membrane potential |
| GO:0050953 | sensory perception of light stimulus |
| GO:0010881 | of cardiac muscle contraction by regulation of the release of sequestered |
| GO:0044295 | axonal growth cone |
| GO:0086091 | regulation of heart rate by cardiac conduction |
| GO:0055117 | regulation of cardiac muscle contraction |
| GO:0045773 | positive regulation of axon extension |
| GO:0097060 | synaptic membrane |
| GO:0030673 | axolemma |
| GO:0086004 | regulation of cardiac muscle cell contraction |
| GO:0098910 | regulation of atrial cardiac muscle cell action potential |
| GO:0086015 | SA node cell action potential |
| GO:0086014 | atrial cardiac muscle cell action potential |
| GO:0098907 | regulation of SA node cell action potential |
| GO:0021782 | glial cell development |
| GO:0035995 | detection of muscle stretch |
| GO:1901631 | positive regulation of presynaptic membrane organization |
| GO:0007608 | sensory perception of smell |
| GO:0007422 | peripheral nervous system development |
| GO:0042826 | histone deacetylase binding |
| GO:0015074 | DNA integration |
| GO:0003964 | RNA-directed DNA polymerase activity |

| | |
|---|---|
| GO:0006364 | rRNA processing |
| GO:0032040 | small-subunit processome |
| GO:0005730 | nucleolus |
| GO:0016513 | core-binding factor complex |
| GO:0000981 | DNA-binding transcription factor activity, RNA polymerase II-specific |
| GO:0005634 | nucleus |
| GO:0015074 | DNA integration |
| GO:0003964 | RNA-directed DNA polymerase activity |
| GO:0010628 | positive regulation of gene expression |
| GO:0043235 | receptor complex |
| GO:0034457 | Mpp10 complex |
| GO:0042567 | insulin-like growth factor ternary complex |
| GO:0048495 | Roundabout binding |
| GO:0050750 | low-density lipoprotein particle receptor binding |
| GO:0043409 | negative regulation of MAPK cascade |
| GO:0043627 | response to estrogen |
| GO:0003081 | regulation of systemic arterial blood pressure by renin-angiotensin |
| GO:0035814 | negative regulation of renal sodium excretion |
| GO:0060252 | positive regulation of glial cell proliferation |
| GO:0009612 | response to mechanical stimulus |
| GO:0038023 | signaling receptor activity |
| GO:0061098 | positive regulation of protein tyrosine kinase activity |
| GO:0006953 | acute-phase response |
| GO:0007154 | cell communication |
| GO:0004978 | corticotropin receptor activity |
| GO:0097720 | calcineurin-mediated signaling |
| GO:0140031 | phosphorylation-dependent protein binding |
| GO:1904209 | positive regulation of chemokine (C-C motif) ligand 2 secretion |
| GO:1904395 | sitive regulation of skeletal muscle acetylcholine-gated channel clusteri |
| GO:0032148 | activation of protein kinase B activity |
| GO:0004879 | nuclear receptor activity |
| GO:0001503 | ossification |
| GO:0015293 | symporter activity |
| GO:0008028 | monocarboxylic acid transmembrane transporter activity |
| GO:0008508 | bile acid:sodium symporter activity |
| GO:0015721 | bile acid and bile salt transport |
| GO:0035725 | sodium ion transmembrane transport |
| GO:0044325 | ion channel binding |
| GO:2001259 | positive regulation of cation channel activity |
| GO:0015718 | monocarboxylic acid transport |
| GO:0042626 | ATPase-coupled transmembrane transporter activity |

| | |
|---|---|
| GO:0005307 | choline:sodium symporter activity |
| GO:0015280 | ligand-gated sodium channel activity |
| GO:0005216 | ion channel activity |
| GO:0034706 | sodium channel complex |
| GO:0015220 | choline transmembrane transporter activity |
| GO:0008273 | calcium, potassium:sodium antiporter activity |
| GO:0006814 | sodium ion transport |
| GO:0005412 | glucose:sodium symporter activity |
| GO:0015871 | choline transport |
| GO:0050891 | multicellular organismal water homeostasis |
| GO:0071805 | potassium ion transmembrane transport |
| GO:0043190 | ATP-binding cassette (ABC) transporter complex |
| GO:1901385 | regulation of voltage-gated calcium channel activity |
| GO:1901018 | positive regulation of potassium ion transmembrane transporter activity |
| GO:0055078 | sodium ion homeostasis |
| GO:0005248 | voltage-gated sodium channel activity |
| GO:0015562 | efflux transmembrane transporter activity |
| GO:0005261 | cation channel activity |
| GO:0008514 | organic anion transmembrane transporter activity |
| GO:0015129 | lactate transmembrane transporter activity |
| GO:0006898 | receptor-mediated endocytosis |
| GO:0070062 | extracellular exosome |
| GO:0038024 | cargo receptor activity |
| GO:0006897 | endocytosis |
| GO:0072659 | protein localization to plasma membrane |
| GO:0005764 | lysosome |
| GO:0070972 | protein localization to endoplasmic reticulum |
| GO:2000008 | regulation of protein localization to cell surface |
| GO:0055085 | transmembrane transport |
| GO:0005319 | lipid transporter activity |
| GO:0006869 | lipid transport |
| GO:0036371 | protein localization to T-tubule |
| GO:0140115 | export across plasma membrane |
| GO:0090370 | negative regulation of cholesterol efflux |

| Odds Ratio | padj | Direction | Category | Subcategory |
|---|---|---|---|---|
| 3.25 | 0.0000 | + | calcium | |
| 10.31 | 0.0004 | + | calcium | |
| 215.35 | 0.0014 | + | calcium | |
| 153.61 | 0.0163 | + | calcium | |
| 153.61 | 0.0163 | + | calcium | |
| 5.99 | 0.0206 | + | calcium | |
| 12.68 | 0.0428 | - | cytoskeleton | adhesion |
| 86.57 | 0.0428 | - | cytoskeleton | adhesion |
| 3.84 | 0.0002 | - | cytoskeleton | ECM |
| 27.62 | 0.0110 | - | cytoskeleton | ECM |
| 2.80 | 0.0177 | - | cytoskeleton | ECM |
| 4.67 | 0.0428 | - | cytoskeleton | ECM |
| 11.44 | 0.0428 | - | cytoskeleton | ECM |
| 8.20 | 0.0289 | - | cytoskeleton | migration |
| 67.86 | 0.0001 | + | cytoskeleton | adhesion |
| 2.59 | 0.0062 | + | cytoskeleton | adhesion |
| 153.61 | 0.0163 | + | cytoskeleton | adhesion |
| 23.92 | 0.0163 | + | cytoskeleton | adhesion |
| 4.20 | 0.0232 | + | cytoskeleton | adhesion |
| 2.99 | 0.0247 | + | cytoskeleton | adhesion |
| 4.10 | 0.0000 | + | cytoskeleton | ECM |
| 3.90 | 0.0000 | + | cytoskeleton | ECM |
| 6.83 | 0.0001 | + | cytoskeleton | ECM |
| 11.69 | 0.0003 | + | cytoskeleton | ECM |
| 5.04 | 0.0003 | + | cytoskeleton | ECM |
| 13.84 | 0.0012 | + | cytoskeleton | ECM |
| 215.35 | 0.0014 | + | cytoskeleton | ECM |
| 6.83 | 0.0030 | + | cytoskeleton | ECM |
| 4.98 | 0.0207 | + | cytoskeleton | ECM |
| 25.20 | 0.0036 | + | cytoskeleton | migration |
| 71.78 | 0.0036 | + | cytoskeleton | migration |
| 35.68 | 0.0000 | + | cytoskeleton | |
| 13.87 | 0.0000 | + | cytoskeleton | |
| 22.09 | 0.0001 | + | cytoskeleton | |
| 4.18 | 0.0001 | + | cytoskeleton | |
| 5.63 | 0.0003 | + | cytoskeleton | |
| 14.05 | 0.0003 | + | cytoskeleton | |
| 26.10 | 0.0007 | + | cytoskeleton | |
| 10.78 | 0.0010 | + | cytoskeleton | |
| 22.62 | 0.0010 | + | cytoskeleton | |

| | | | | |
|---|---|---|---|---|
| 7.62 | 0.0018 | + | cytoskeleton | |
| 4.01 | 0.0018 | + | cytoskeleton | |
| 71.78 | 0.0036 | + | cytoskeleton | |
| 71.78 | 0.0036 | + | cytoskeleton | |
| 9.79 | 0.0038 | + | cytoskeleton | |
| 5.39 | 0.0044 | + | cytoskeleton | |
| 6.02 | 0.0114 | + | cytoskeleton | |
| 30.76 | 0.0125 | + | cytoskeleton | |
| 153.61 | 0.0163 | + | cytoskeleton | |
| 9.55 | 0.0293 | + | cytoskeleton | |
| 16.56 | 0.0303 | + | cytoskeleton | |
| 51.20 | 0.0334 | + | cytoskeleton | |
| 2.50 | 0.0372 | + | cytoskeleton | |
| 3.58 | 0.0177 | - | development | |
| 19.17 | 0.0468 | - | development | |
| 9.86 | 0.0014 | + | development | |
| 10.85 | 0.0029 | + | development | |
| 10.29 | 0.0035 | + | development | |
| 43.07 | 0.0075 | + | development | |
| 14.59 | 0.0125 | + | development | |
| 153.61 | 0.0163 | + | development | |
| 153.61 | 0.0163 | + | development | |
| 23.92 | 0.0163 | + | development | |
| 5.35 | 0.0299 | + | development | |
| 3.39 | 0.0351 | + | development | |
| 259.72 | 0.0295 | - | immune | stress |
| 86.57 | 0.0428 | - | immune | |
| 86.57 | 0.0428 | - | immune | |
| 153.61 | 0.0163 | + | immune | stress |
| 14.48 | 0.0000 | + | immune | |
| 19.95 | 0.0014 | + | immune | |
| 5.94 | 0.0029 | + | immune | |
| 71.78 | 0.0036 | + | immune | |
| 19.57 | 0.0227 | + | immune | |
| 19.57 | 0.0227 | + | immune | |
| 51.20 | 0.0334 | + | immune | |
| 7.92 | 0.0412 | + | immune | |
| 7.92 | 0.0412 | + | immune | |
| 2.95 | 0.0000 | + | membrane | |
| 2.64 | 0.0000 | + | membrane | |
| 4.97 | 0.0000 | + | membrane | |

| | | | | |
|---|---|---|---|---|
| 35.10 | 0.0000 | + | membrane | |
| 5.10 | 0.0000 | + | membrane | |
| 2.69 | 0.0000 | + | membrane | |
| 10.35 | 0.0000 | + | membrane | |
| 2.31 | 0.0001 | + | membrane | |
| 4.79 | 0.0009 | + | membrane | |
| 13.84 | 0.0012 | + | membrane | |
| 17.85 | 0.0019 | + | membrane | |
| 5.12 | 0.0033 | + | membrane | |
| 6.80 | 0.0152 | + | membrane | |
| 153.61 | 0.0163 | + | membrane | |
| 5.20 | 0.0169 | + | membrane | |
| 6.65 | 0.0303 | + | membrane | |
| 3.04 | 0.0010 | - | metabolic | prosthetic |
| 18.78 | 0.0253 | - | metabolic | regulation |
| 61.94 | 0.0000 | - | metabolic | |
| 8.27 | 0.0002 | - | metabolic | |
| 25.00 | 0.0021 | - | metabolic | |
| 10.84 | 0.0289 | - | metabolic | |
| 259.72 | 0.0295 | - | metabolic | |
| 7.47 | 0.0295 | - | metabolic | |
| 15.14 | 0.0295 | - | metabolic | |
| 28.03 | 0.0373 | - | metabolic | |
| 11.44 | 0.0428 | - | metabolic | |
| 86.57 | 0.0428 | - | metabolic | |
| 19.17 | 0.0468 | - | metabolic | |
| 2.59 | 0.0001 | + | metabolic | prosthetic |
| 16.98 | 0.0000 | + | metabolic | prosthetic |
| 92.42 | 0.0003 | + | metabolic | regulation |
| 92.42 | 0.0003 | + | metabolic | regulation |
| 4.29 | 0.0031 | + | metabolic | regulation |
| 8.54 | 0.0066 | + | metabolic | regulation |
| 30.76 | 0.0125 | + | metabolic | regulation |
| 19.57 | 0.0227 | + | metabolic | regulation |
| 19.57 | 0.0227 | + | metabolic | regulation |
| 5.07 | 0.0334 | + | metabolic | regulation |
| 6.16 | 0.0350 | + | metabolic | regulation |
| 5.95 | 0.0385 | + | metabolic | regulation |
| 12.66 | 0.0445 | + | metabolic | regulation |
| 3.89 | 0.0486 | + | metabolic | regulation |
| 21.32 | 0.0049 | + | metabolic | |

| | | | | |
|---|---|---|---|---|
| 71.78 | 0.0036 | + | metabolic | |
| 14.59 | 0.0125 | + | metabolic | |
| 14.35 | 0.0351 | + | metabolic | |
| 92.42 | 0.0003 | + | metabolic | |
| 92.42 | 0.0003 | + | metabolic | |
| 55.45 | 0.0008 | + | metabolic | |
| 5.71 | 0.0009 | + | metabolic | |
| 4.49 | 0.0014 | + | metabolic | |
| 16.15 | 0.0026 | + | metabolic | |
| 7.85 | 0.0036 | + | metabolic | |
| 71.78 | 0.0036 | + | metabolic | |
| 21.32 | 0.0049 | + | metabolic | |
| 18.48 | 0.0071 | + | metabolic | |
| 43.07 | 0.0075 | + | metabolic | |
| 43.07 | 0.0075 | + | metabolic | |
| 14.59 | 0.0125 | + | metabolic | |
| 30.76 | 0.0125 | + | metabolic | |
| 30.76 | 0.0125 | + | metabolic | |
| 5.58 | 0.0152 | + | metabolic | |
| 8.69 | 0.0163 | + | metabolic | |
| 6.37 | 0.0163 | + | metabolic | |
| 153.61 | 0.0163 | + | metabolic | |
| 153.61 | 0.0163 | + | metabolic | |
| 153.61 | 0.0163 | + | metabolic | |
| 153.61 | 0.0163 | + | metabolic | |
| 153.61 | 0.0163 | + | metabolic | |
| 153.61 | 0.0163 | + | metabolic | |
| 23.92 | 0.0163 | + | metabolic | |
| 5.81 | 0.0227 | + | metabolic | |
| 19.57 | 0.0227 | + | metabolic | |
| 10.26 | 0.0239 | + | metabolic | |
| 16.56 | 0.0303 | + | metabolic | |
| 51.20 | 0.0334 | + | metabolic | |
| 51.20 | 0.0334 | + | metabolic | |
| 51.20 | 0.0334 | + | metabolic | |
| 51.20 | 0.0334 | + | metabolic | |
| 14.35 | 0.0351 | + | metabolic | |
| 7.92 | 0.0412 | + | metabolic | |
| 5.74 | 0.0430 | + | metabolic | |
| 12.66 | 0.0445 | + | metabolic | |
| 1.43 | 0.0169 | + | negligible | cytoplasm |

| | | | | |
|---|---|---|---|---|
| 86.57 | 0.0428 | - | neural | |
| 4.39 | 0.0440 | - | neural | |
| 16.30 | 0.0097 | + | neural | |
| 5.75 | 0.0001 | + | neural | |
| 8.04 | 0.0003 | + | neural | |
| 10.78 | 0.0010 | + | neural | |
| 22.62 | 0.0010 | + | neural | |
| 30.80 | 0.0024 | + | neural | |
| 71.78 | 0.0036 | + | neural | |
| 21.32 | 0.0049 | + | neural | |
| 5.91 | 0.0057 | + | neural | |
| 43.07 | 0.0075 | + | neural | |
| 16.30 | 0.0097 | + | neural | |
| 10.28 | 0.0099 | + | neural | |
| 2.56 | 0.0152 | + | neural | |
| 153.61 | 0.0163 | + | neural | |
| 153.61 | 0.0163 | + | neural | |
| 153.61 | 0.0163 | + | neural | |
| 153.61 | 0.0163 | + | neural | |
| 153.61 | 0.0163 | + | neural | |
| 23.92 | 0.0163 | + | neural | |
| 23.92 | 0.0163 | + | neural | |
| 11.08 | 0.0207 | + | neural | |
| 19.57 | 0.0227 | + | neural | |
| 10.26 | 0.0239 | + | neural | |
| 9.55 | 0.0293 | + | neural | |
| 8.94 | 0.0334 | + | neural | |
| 8.94 | 0.0334 | + | neural | |
| 51.20 | 0.0334 | + | neural | |
| 51.20 | 0.0334 | + | neural | |
| 51.20 | 0.0334 | + | neural | |
| 51.20 | 0.0334 | + | neural | |
| 51.20 | 0.0334 | + | neural | |
| 51.20 | 0.0334 | + | neural | |
| 51.20 | 0.0334 | + | neural | |
| 51.20 | 0.0334 | + | neural | |
| 7.92 | 0.0412 | + | neural | |
| 12.66 | 0.0445 | + | neural | |
| 8.33 | 0.0428 | - | nucleic acid | chromatin |
| 15.83 | 0.0021 | - | nucleic acid | recomb |
| 18.25 | 0.0001 | - | nucleic acid | replication |

| | | | | |
|---|---|---|---|---|
| 5.74 | 0.0375 | - | nucleic acid | ribosome |
| 12.68 | 0.0428 | - | nucleic acid | ribosome |
| 2.59 | 0.0428 | - | nucleic acid | ribosome |
| 86.57 | 0.0428 | - | nucleic acid | transcription |
| 4.00 | 0.0289 | - | nucleic acid | transcription |
| 1.60 | 0.0428 | - | nucleic acid | |
| 11.31 | 0.0009 | + | nucleic acid | recomb |
| 7.57 | 0.0104 | + | nucleic acid | replication |
| 3.03 | 0.0327 | + | nucleic acid | transcription |
| 5.58 | 0.0177 | - | signaling | |
| 259.72 | 0.0295 | - | signaling | |
| 259.72 | 0.0295 | - | signaling | |
| 21.43 | 0.0428 | - | signaling | |
| 21.43 | 0.0428 | - | signaling | |
| 19.17 | 0.0468 | - | signaling | |
| 11.89 | 0.0007 | + | signaling | |
| 55.45 | 0.0008 | + | signaling | |
| 153.61 | 0.0163 | + | signaling | |
| 19.57 | 0.0227 | + | signaling | |
| 10.26 | 0.0239 | + | signaling | |
| 3.60 | 0.0296 | + | signaling | |
| 16.56 | 0.0303 | + | signaling | |
| 16.56 | 0.0303 | + | signaling | |
| 16.56 | 0.0303 | + | signaling | |
| 51.20 | 0.0334 | + | signaling | |
| 51.20 | 0.0334 | + | signaling | |
| 51.20 | 0.0334 | + | signaling | |
| 51.20 | 0.0334 | + | signaling | |
| 51.20 | 0.0334 | + | signaling | |
| 12.66 | 0.0445 | + | signaling | |
| 7.49 | 0.0474 | + | signaling | |
| 10.91 | 0.0450 | - | skeleton | |
| 9.24 | 0.0000 | + | transport | channel/pump |
| 19.00 | 0.0000 | + | transport | channel/pump |
| 36.50 | 0.0001 | + | transport | channel/pump |
| 20.17 | 0.0001 | + | transport | channel/pump |
| 13.25 | 0.0004 | + | transport | channel/pump |
| 8.06 | 0.0006 | + | transport | channel/pump |
| 39.60 | 0.0014 | + | transport | channel/pump |
| 17.85 | 0.0019 | + | transport | channel/pump |
| 6.49 | 0.0036 | + | transport | channel/pump |

| | | | | |
|---|---|---|---|---|
| 71.78 | 0.0036 | + | transport | channel/pump |
| 9.69 | 0.0121 | + | transport | channel/pump |
| 14.59 | 0.0125 | + | transport | channel/pump |
| 30.76 | 0.0125 | + | transport | channel/pump |
| 23.92 | 0.0163 | + | transport | channel/pump |
| 23.92 | 0.0163 | + | transport | channel/pump |
| 5.99 | 0.0206 | + | transport | channel/pump |
| 19.57 | 0.0227 | + | transport | channel/pump |
| 19.57 | 0.0227 | + | transport | channel/pump |
| 19.57 | 0.0227 | + | transport | channel/pump |
| 5.49 | 0.0271 | + | transport | channel/pump |
| 51.20 | 0.0334 | + | transport | channel/pump |
| 51.20 | 0.0334 | + | transport | channel/pump |
| 51.20 | 0.0334 | + | transport | channel/pump |
| 14.35 | 0.0351 | + | transport | channel/pump |
| 14.35 | 0.0351 | + | transport | channel/pump |
| 14.35 | 0.0351 | + | transport | channel/pump |
| 8.40 | 0.0351 | + | transport | channel/pump |
| 12.66 | 0.0445 | + | transport | channel/pump |
| 12.66 | 0.0445 | + | transport | channel/pump |
| 8.46 | 0.0002 | + | transport | vesicular |
| 2.94 | 0.0014 | + | transport | vesicular |
| 16.15 | 0.0026 | + | transport | vesicular |
| 3.71 | 0.0163 | + | transport | vesicular |
| 5.20 | 0.0169 | + | transport | vesicular |
| 2.86 | 0.0238 | + | transport | vesicular |
| 51.20 | 0.0334 | + | transport | vesicular |
| 51.20 | 0.0334 | + | transport | vesicular |
| 6.06 | 0.0026 | + | transport | |
| 18.48 | 0.0071 | + | transport | |
| 7.87 | 0.0090 | + | transport | |
| 153.61 | 0.0163 | + | transport | |
| 23.92 | 0.0163 | + | transport | |
| 51.20 | 0.0334 | + | transport | |
